# Invasive species modulate the structure and stability of a multilayer mutualistic network

**DOI:** 10.1101/2022.12.25.521894

**Authors:** Agustin Vitali, Sofía Ruiz-Suarez, Diego P. Vázquez, Matthias Schleuning, Mariano A. Rodríguez-Cabal, Yamila Sasal, Shai Pilosof

## Abstract

Species interactions are the backbone of the structure and dynamics of communities. The extensive research into the link between structure and stability has been primarily theoretical and focused on monotrophic networks. Therefore, how the disruption of multitrophic interactions alters communities’ response to perturbations in nature remains an open question. Here, we explored how non-native ungulates affect pollination-seed dispersal multilayer networks in Patagonia, Argentina. Ungulates disrupt a hummingbird-mistletoe-marsupial keystone interaction, which alters community composition. We calculated interlayer connectivity, modularity, and species’ roles in connecting modules for intact vs. invaded networks. To link structural changes to stability, we quantified network tolerance to a single random species removal (disturbance propagation) and sequential species removal (robustness) using a stochastic coextinction model. Non-native ungulates reduced the connectivity between pollination and seed dispersal and produced fewer modules with a skewed size distribution. Moreover, species shifted their structural role, primarily from connectors to peripherals, thereby fragmenting the network by reducing the “bridges” among modules. These structural changes altered the dynamics of cascading effects in the community, increasing disturbance propagation and reducing network robustness. Our results highlight the importance of understanding the mechanisms that alter the structure and subsequent stability of multitrophic communities in nature.

## Introduction

Ecological interactions are the backbone of the structure and dynamics of communities [1,2]. Therefore, understanding how the disruption of species interactions affects community stability is fundamental to coping with anthropogenic effects. The vast research on the link between structure and stability has focused on monotrophic networks (e.g., food webs) [3, 4]. Nevertheless, in natural communities, species are often involved in multitrophic interactions. For instance, flowering plants may depend on both pollination and seed dispersal. The few works on structure-stability in multitrophic communities focused on mutualistic-antagonistic interactions, and used simulated data and theoretical modeling [5–8]. Although critical to developing such theory, studies that directly compare the structure and stability in disturbed and undisturbed natural sites are still needed. This is particularly important due to the current rates of invasive species and consequent biodiversity loss [9]. Hence, the effect of invasive species on the response of multitrophic communities’ to perturbations in nature remains an open question.

The link between the structure and stability of ecological communities is well studied in networks with a single kind of interaction (e.g., pollination or predation) [10, 11]. For instance, a modular structure—in which species within modules interact more frequently than with species from other modules—increases stability because perturbations tend to be locked within modules before spreading to other modules [12]. To date, however, modularity has been rarely studied in multitrophic networks. Moreover, species play different structural roles according to the distribution of their interactions across partners within and among modules, influencing network cohesion [13], and hence, the dynamics of cascading effects [14, 15]. For example, a high proportion of species that connect modules (i.e., connector species) may favor disturbance propagation across modules. However, the same species can play different structural roles in different trophic interactions. Furthermore, when multiple types of interactions are considered, the connectivity between different trophic groups could also affect the propagation of disturbances. For instance, the robustness of parrot communities increased when both antagonistic and mutualistic interactions with plants were considered [7].

Early works have emphasized the importance of keystone species to the stability of ecological communities. Analogously to keystone species, keystone interactions are those that determine structural and functional properties of communities [16, 17]. For instance, in the temperate forest of Patagonia, a multitrophic keystone interaction between a hummingbird, a mistletoe and a seed disperser marsupial was identified (Fig. 1) [18, 19]. The mistletoe provides a unique nectar resource for the generalist hummingbird during winter, supporting resident populations in the forest [20, 21]. Its fruits allow the marsupial to increase its abundance and hence sustain the dispersal of other fleshy fruited species [22, 23]. Non-native ungulates (deer and cattle; Methods) disrupt this keystone interaction via herbivory on the main host of the mistletoe, *Aristotelia chilensis*, and by causing changes to the vegetation structure. This disruption reduces the complexity of pollination and seed dispersal networks [19].

**Fig. 1:**
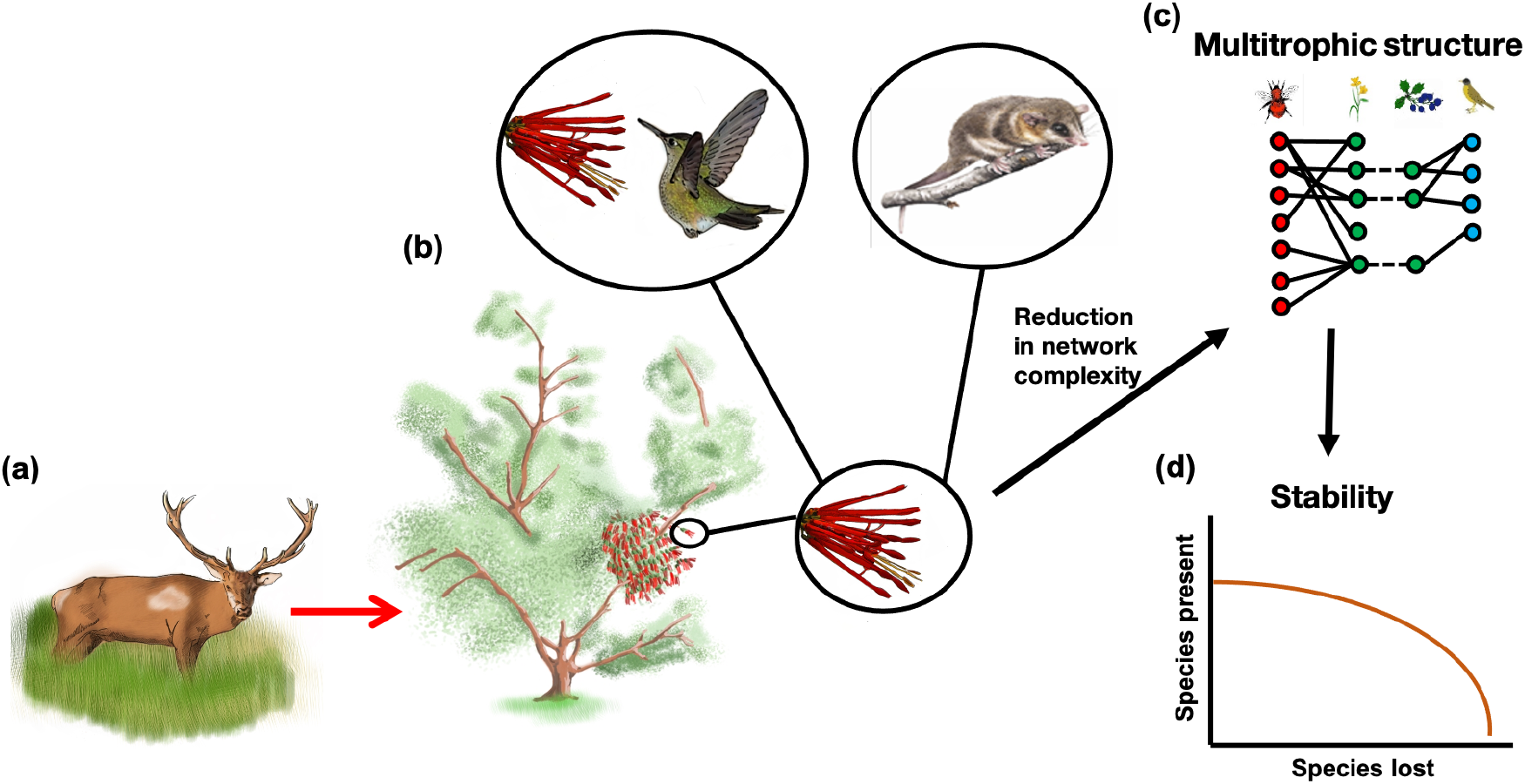
Study system and hypothesis. The mistletoe (*Tristerix corymbosus*) grows on its host *Aristotelia chilensis* and provides nectar to the migratory hummingbird (*Sephanoides sephaniodes*) and fruits for the seed disperser marsupial (*Dromiciops gliroides*), sustaining their populations. The hummingbird pollinates almost 20% of woody plants in summer [21, 25] and the marsupial disperses up to 58% of fleshy fruited species, providing a crucial service for plants with large fruits that are not dispersed by native birds [26, 27]. In addition, the marsupial is the only seed disperser of the mistletoe, facilitating the seeds’ placement on branches of an appropriate host [26]. **(a)** Disruption by non-native ungulates (red arrow) leads to the extinction of the hummingbird-mistletoe-marsupial keystone interaction by consuming the main host of the mistletoe, and via local extinction of the marsupial **(b)**. The disruption of this keystone interaction produces cascading effects that reduce the number of species and interactions in pollination and seed dispersal networks [19]. However, how the complete multitrophic structure is altered remains unknown. **(c) and (d)** We hypothesized that non-native ungulates would alter the structure of the pollinator-plantseed disperser multilayer network, affecting the stability of the community. Panels (a) and (b) were adapted from [18].

While it is necessary to consider keystone interactions when studying the link between structure and stability [16, 24], the effect of their disruption is not well understood, particularly for multitrophic networks. Here, we leveraged the Patagonian hummingbird-mistletoe-marsupial keystone interaction and its disruption by an ongoing ungulate invasion to address this gap (Fig. 1). An ideal framework to link multiple trophic interaction networks is with multilayer networks. While ecological multilayer networks are increasingly used, how their structure relates to stability is still grossly understudied. Our model system and multilayer network analysis allows us to study broad questions on how disturbances, such as invasive species, and the subsequent loss of keystone interactions affect the structure and stability of multitrophic networks.

## Results

The study was conducted in the Llao Llao Municipal Reserve and Nahuel Huapi National Park in Patagonia, Argentina. Currently, 56% of the area of Nahuel Huapi National Park is occupied by the non-native ungulates. We built ‘invaded’ and ‘intact’ multilayer networks (Methods). Each network included four components as follows (Fig. S1) [28, 29].

1. Two layers representing pollination (*α*) and seed dispersal (*β*) interactions.
2. Three sets of nodes representing plant (plants can be present in both layers), pollinator (in layer *α*), and seed disperser (in layer *β*) species.
3. Intralayer weighted links connecting plant *j* with pollinator *i*, defined as 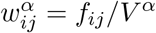, where *f*_*ij*_ denote the number of visits of *i* to *j* and *V* ^*α*^ is the total number of visits in layer *α*. Similarly, intralayer weighted links connecting plant *j* with seed disperser *k*, analogously defined as 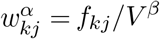.
4. Interlayer weighted links connecting each plant species with itself when it was both pollinated and dispersed by animals. Biologically, interlayer links encode the extent to which plants mediate the biomass or energy flow between pollination and seed dispersal. We therefore defined an indirect link as a walk from a node in one layer to a node in another via a plant (Fig. S2). We then calculated interlayer link weight as 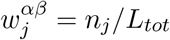, where *n*_*j*_ is the total number of indirect links between any pollinator and seed disperser species mediated by the plant species *j*, and *L*_*tot*_ is the total number of indirect links connecting all pollinator and seed disperser species.

Note that the weight of both intralayer and interlayer links ranged between 0 and 1. This places the two link types on the same scale, ensuring that network properties are not a-priori biased towards any of these [28, 30].

We hypothesized that non-native ungulates would alter the structure of the network. Previous studies found that non-native ungulates disrupt mutualistic interactions [31] and reduce the complexity of pollination and seed dispersal networks [19]. In our networks, the invaded network contained less species than the intact (24 vs 37 plants, 67 vs 95 pollinators; Table S6). We therefore expected a higher connectivity between the trophic levels and a greater number of modules in the intact than in the invaded network. In addition, we expected species to change their structural role between the intact and invaded networks, as has been recorded for other disturbances [32]. We tested this hypothesis by measuring connectivity between trophic levels, modularity, and the structural role of species.

### Invasive ungulates altered network connectivity

We conducted two analyses to estimate the connectivity between pollination and seed dispersal. First, we used generalized linear mixed models (GLMMs) to test whether the presence of plant species *j* connecting both mutualisms and its interlayer connectivity (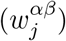) differed between the intact and invaded networks. Non-native ungulates reduced the connectivity between pollination and seed dispersal: The presence of plant species connecting pollination and seed dispersal mutualism was almost double in the intact (8 species) than in the invaded (5 species) network (GLMM, *z* = 2.617, *P <* 0.01). Nevertheless, they represented almost the same proportion of species (22% and 21% of total species, respectively). In addition, we did not find differences in plants’ interlayer link weights between the intact and invaded network (GLMM, *z* = 0.358, *P* = 0.721), suggesting that the ability of plants to connect between trophic levels is similar regardless of the presence of non-native ungulates.

Second, we assessed if non-native ungulates change the connectivity between pollinator and seed disperser species. To do this, we built a binary matrix for the intact and invaded networks, encoding the existence of at least one indirect link between both trophic groups and calculated the proportion of indirect links per pollinator and seed disperser species (Fig. S3). Then, we performed a GLMM for each trophic group with the proportion of indirect links per species as a response variable and network types as an explanatory variable. The proportion of indirect links per pollinator (GLMM, *z* = 5.642, *P <* 0.001) and seed disperser species (GLMM, *z* = 0.79, *P* = 0.42) were greater in the intact network, indicating a stronger connection between both trophic groups. In the intact network, pollinators and seed dispersers had 56% and 59% more indirect links (*pol* = 2.75 *±* 0.14; *disp* = 34.25 *±* 6.8) than in the invaded network (*pol* = 1.76 *±* 0.17; *disp* = 21.6 *±* 6.7). Moreover, the identity of indirect links between pollinators and seed dispersers was highly dissimilar between network types (Jaccard dissimilarity index = 0.82), indicating that the differences between these networks were not solely numeric.

### Invasive ungulates reduced module number and altered the structural role of species

The difference in connectivity between the two networks implies that they may also differ in the degree of modularity–a highly relevant property for stability. For each type of network, we identified groups of tightly connected species with a modularity analysis using Infomap [33]. We also assessed if the observed modularity was a result of random processes. The observed number of modules was different than shuffled networks in both networks (*P* < 0.001; Fig. S4), indicating that their modular structure was non-random. In both the intact and invaded networks, five (28%) and three (27%) modules contained species from the three species groups, respectively. However, the intact network had 63% more modules than the invaded network (18 vs 11) (Fig. 2).

**Fig. 2:**
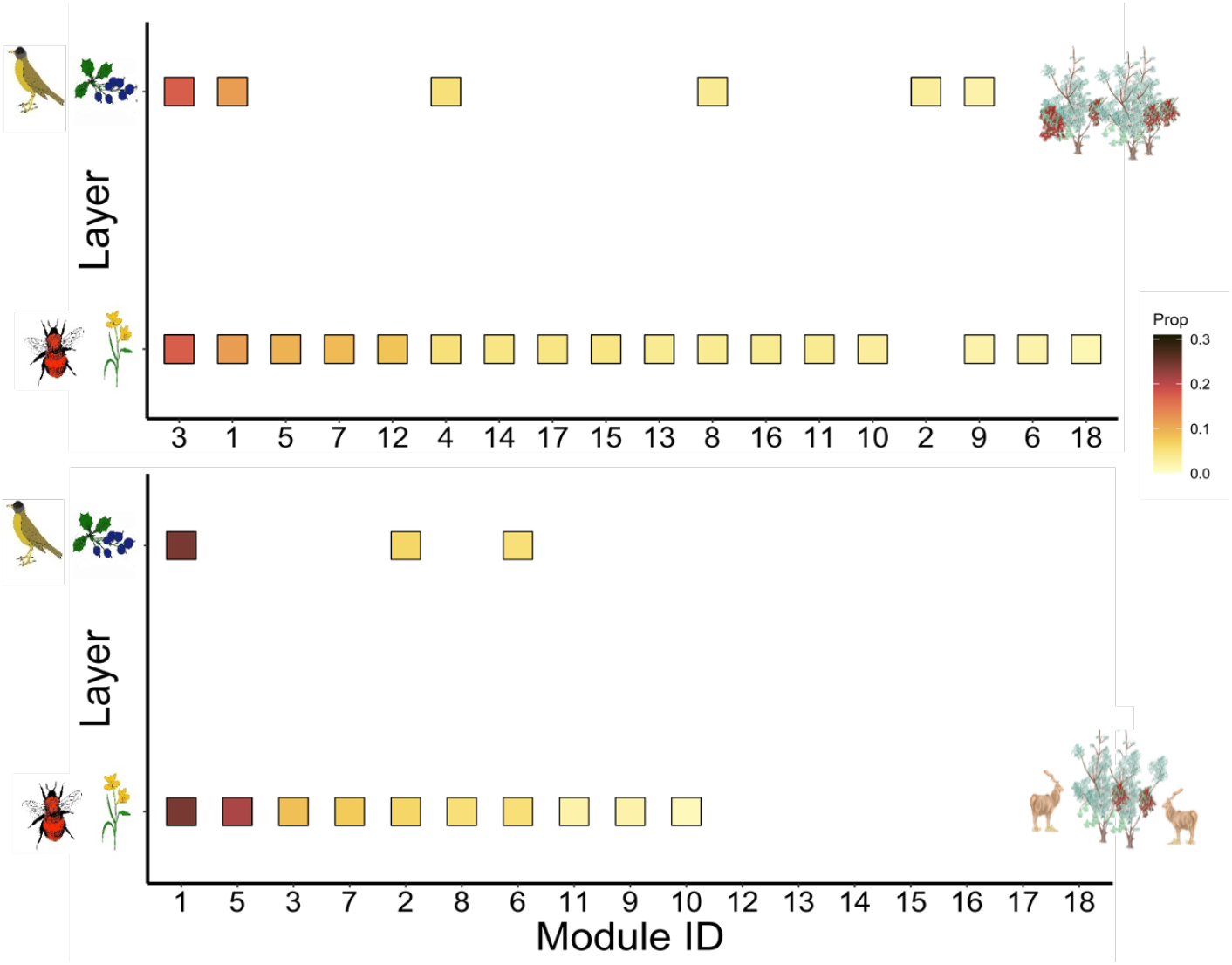
Modular structure differs between the networks. Module-layer combinations are shown for the intact (upper panel) and invaded (lower panel) networks. Modules can span layers and each square represents the occurrence of a specific module in a layer. While most modules were limited to a single layer, a few spanned both layers (e.g., module 1 in both networks). Square color depicts the proportion of species in the network that were assigned to the module. Modules are ordered from higher to lower size proportion (from left to right). Module IDs are assigned randomly and independently for the two networks (e.g., module 1 in the intact network is not the same as in the invaded).

Nevertheless, difference in module number alone is not enough to provide stability. For perturbation to spread between modules it is also crucial to test how the modules are connected. We assessed network fragmentation by estimating the structural role of species. Most species, and mainly pollinators, were classified as peripherals in both networks but the percentage of peripheral species was lower in the intact than in the invaded network (57% and 71%, respectively; Fig. S6). In contrast, the percentage of connector species was higher in the intact (41%) than in the invaded network (27%), which indicates less connection among modules and, therefore, a more fragmented network in presence of non-native ungulates. Specifically, two seed disperser species acted as connectors (*D. gliroides* and the generalist bird *Elaenia albiceps*) in the intact network but no seed dispersers were connectors in the invaded network. Only plants acted as module hubs in both the intact and invaded networks. However, we detected network hubs only in the intact network: the plants *Alstromeria aurea* and *Schinus patagonicus*.

Ungulate invasion changed the structural role of 46% of plants in the pollination layer, 35% of pollinators, and 33% of seed dispersers. However, plants in the seed dispersal layer did not change their structural role. While most species (73%) changed their role from connector to peripheral, a few shifted from network hub to connector (4%) or to module hub (4%) (Fig. 3).

**Fig. 3:**
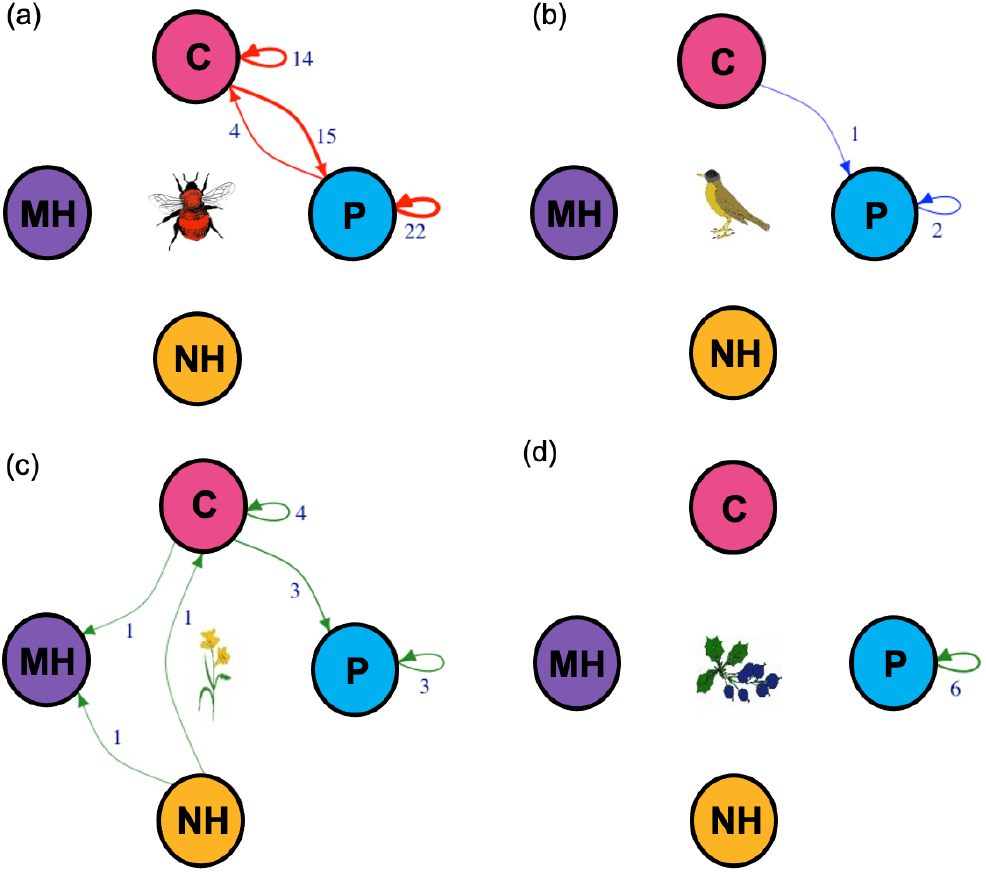
Changes in species structural roles. Circles indicate species’ structural roles: P = peripheral (light blue), C = connector (pink), MH = module hub (purple), NH = network hub (orange). Changes in the structural role are represented by arrows. Origin and end of arrows represent the role of species in the intact (“from”) and in the invaded (“to”) network, respectively. Self-loop indicates no change in the structural role. Values in each arrow indicate the number of species that experienced (or not) the change in their structural role. Panels represent different trophic groups: **(a)** pollinators, **(b)** seed dispersers, **(c)** plants in the pollination layer, and **(d)** plants in the seed dispersal layer. Only specie shared between network types are included in the calculations and figure.

### Invasive ungulates reduced network stability

We hypothesized that by disrupting the network structure, non-native ungulates would alter the dynamics of multitrophic cascading effects and, therefore, network stability (H2). We expected a lower tolerance to the removal of one randomly selected species (i.e., higher disturbance propagation) and to the sequential removal of species (i.e., lower network robustness) in the invaded network.

To capture different aspects of stability in the intact and invaded networks, we evaluated its tolerance to single and sequential species removal [34–36]. The former type of extinction allows us to estimate the propagation after a particular disturbance event (e.g., extinction of a species), while the latter estimates the tolerance of the network to continued disturbances before collapsing (also typically called robustness). To simulate both types of extinctions, we used a stochastic coextinction model, which we modified to include three different trophic groups [35]. This method is optimal to evaluate extinction cascades because it considers the intrinsic demographic dependence of each species on mutualism and incorporates the mutual dependence of each species on its mutualistic partners [35] (Methods and Supplementary Information). To link modularity and species roles to stability, we removed species according to species roles, rather than by degree as is commonly done.

The propagation of disturbances through the community was affected by the statistical interaction between the network types and the structural role of the removed species (GLMM, 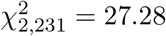, *P* <, Fig. 4a). In the intact network, the removal of network hubs resulted in a 6.9 times higher species extinction than the removal of peripheral species (post-hoc test, *z* = *-*8.700, *P* < 0.01). Similarly, the removal of module hubs produced a 4.7 higher percentage of species extinction than removal of peripheral species in the invaded network (post-hoc test, *z* = *-*4.867, *P* < 0.01). In addition, on average, the propagation of disturbances produced 1.5 times higher percentages of species extinction after the removal of a single species in the invaded than in the intact network. The robustness of communities was affected by the interaction between network types and the order of species removal (GLM, 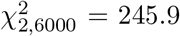, *P* < 0.001, Fig. 4b). The intact network was more robust than the invaded network in the least-to-most (order: peripheral, connector, module hub, network hub) (post-hoc test, t ratio = 23.16, *P* < 0.001) and most-to-least removal orders (post-hoc test, t ratio = 5.16, *P* < 0.001), but not when removing species at random (post-hoc test, t ratio = 0.67, *P* = 0.984).

**Fig. 4:**
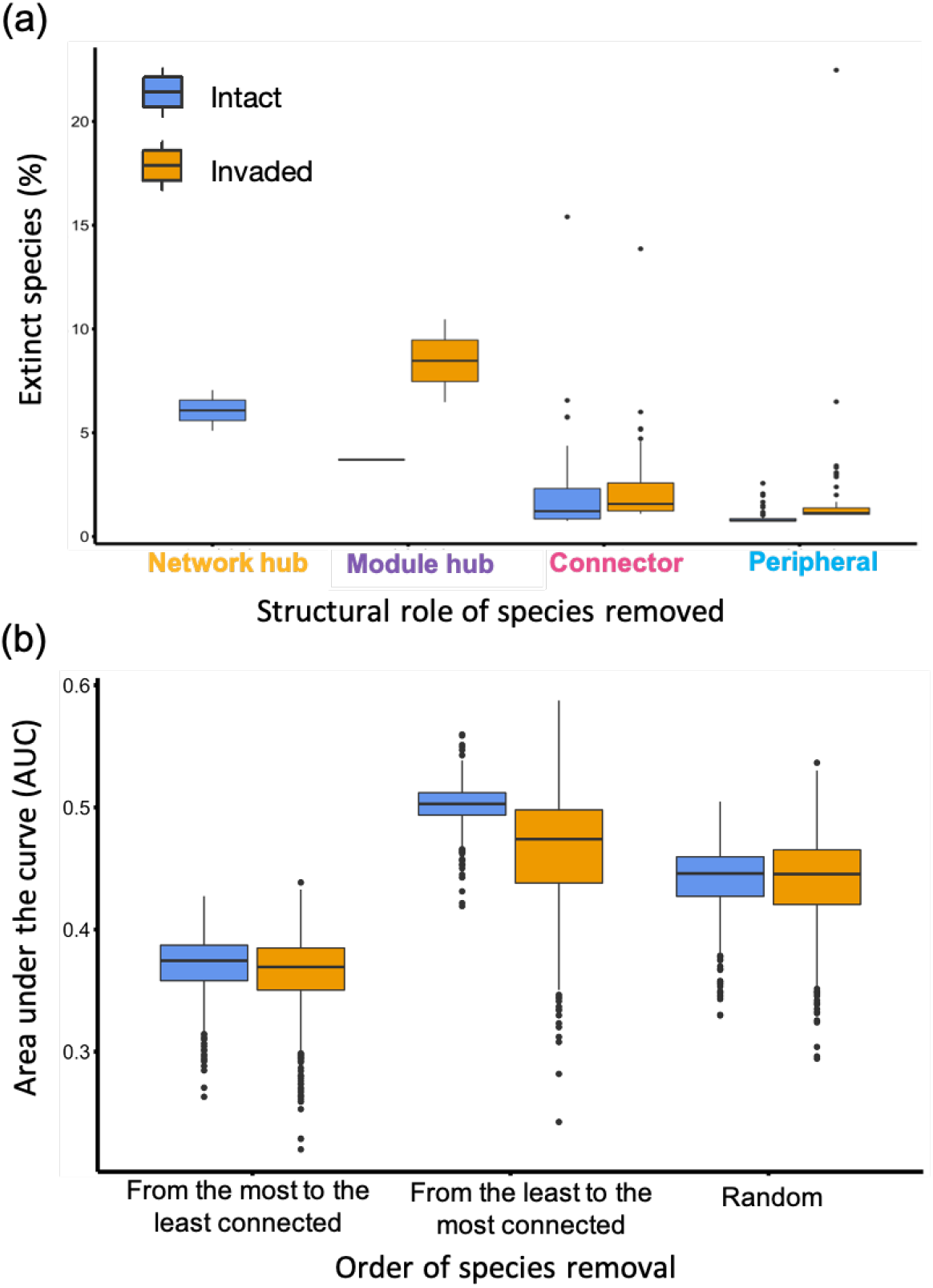
Multitrophic network structure affects stability. Propagation of disturbances was higher (a) and robustness lower (b) in the invaded network. (a) The propagation of disturbance was measured as the proportion of extinct species calculated at equilibrium following the removal of a single species (*E*, y axis). There were seven combinations between structural roles of species removed and network types because there were no network hubs in the invaded network. *E* was affected by the statistical interaction between the network types and the structural role of the removed species. (b) Robustness was measured as the area under an extinction curve (AUC). Most-to-least removal order: network hub, module hub, connector, peripheral. Least-to-most: the opposite order of most-to-least).

## Discussion

Understanding how disturbances reshape the structure of communities and alter their ability to cope with perturbations is essential due to the growing rates of species invasion and biodiversity loss [9, 37]. We showed that the loss of a keystone interaction by invasive ungulates reduces the stability of multitophic mutualistic communities by altering indirect connectivity, the modular structure and associated species roles.

Relatively few studies have investigated structure and stability in multitrophic communities. These studies have focused on disentangling how the topology of each trophic interaction separately affects the stability of the whole community [6,8,38]. For example, the stabilizing mechanisms in antagonistic (modularity) and mutualistic (nestedness) networks have a weak effect when considering both interactions [6]. Despite the relevance of these findings to nature, most studies evaluated mutualistic and antagonist interactions (mainly pollinator-plant and plant-herbivory) using simulated data [6, 8, 38]. One study that used empirical data showed that the combination of properties enhancing robustness in antagonistic (modularity) and mutualistic (nestedness) networks led to an increase in the robustness of a parrot community when including both types of ecological interactions [7].

Here, we studied for the first time how invasive species affect the link between structure and stability in a multitrophic mutualistic community. Our empirical approach allowed us to control for the invasion scenario. A previous study found that logging disturbance produced correlated effects seed dispersal and pollination. However, the indirect links between these two mutualisms and the subsequent effects on stability were not studied [39]. The analytical multilayer approach and dedicated algorithms we applied allowed us to explicitly quantify the propagation of disturbances from one interaction type to another. We encourage future studies to use multilayer networks to understand the response of multitrophic communities to perturbations.

The network had a marked signature of structural reorganization following invasion. Non-native ungulates reduced the connectivity between pollination and seed dispersal mutualisms and caused link turnover. Link turnover could result from species turnover and/or interaction rewiring between animals and plants in the community [40], two well-documented mechanisms to cope with fluctuation in resource availability in animals [41,42]. We further observed a lower number of modules with a skewed size distribution in the invaded network. This could result from the local extinction of species and interactions because networks with a smaller size tend to contain a lower number of modules [43, 44]. Local extinction can happen due to the disruption of the keystone interaction, and the subsequent changes to the vegetation [19, 45]. At the node level, species changed their structural role, primarily from connectors to peripherals. These shifts fragmented the network by reducing the “bridges” among modules [13]. Previous studies also demonstrated the structural role of species change in response to disturbances, such as fire and habitat loss [32, 46].

Changes to network structure altered the dynamics of cascading effects, increasing the propagation of disturbances and reducing the robustness of the invaded network. Specifically, disturbance propagation was more significant in the presence of non-native ungulates. A plausible explanation is the presence of a few large and isolated modules containing many peripheral species. Such organization concentrates propagation within modules and likely produces a high percentage of species extinction after removing a module hub [47,48], collapsing the whole module. Conversely, in the intact network, the large number of connected modules containing a low proportion of species allows propagation to spread across the network, limiting the collapse of entire modules. By controlling for network size, we further found that the high number of species in the intact network reduced the propagation of disturbances. A negative correlation between network size and disturbance propagation has been found in mutualistic networks [49]. As with propagation, the intact network was more robust to the sequential species removal. This result is consistent with literature showing that a higher number of modules slows down cascading effects through the network, increasing the tolerance of the network to collapse [15, 50]. In particular, the strong robustness to the sequential removal of peripheral species in the intact network indicates a higher tolerance of the community to common disturbance scenarios such as habitat fragmentation and climate change, which primarily affect rare and specialized species [51].

Our study has two main limitations. First, we uncovered the impacts of invasive species on the structure and dynamics of communities 100 years after the introduction of invasive species. Nevertheless, invasion dynamics encompass different stages (e.g., establishment and spread) in which exotic species interact differently with species in the new community before becoming invasive [52, 53]. Hence, we encourage future studies to focus on how exotic species reshape structure and stability across different invasion stages using temporal networks. Second, the stochastic extinction model does not incorporate all ecological processes occurring in natural systems [54, 55]. For example, interaction rewiring was not included. Nevertheless, we increased the biological realism by considering the extent of species dependence on the mutualistic interactions using data on species diet [35]. A relevant open question is how different trophic groups respond to extinction cascades. Our preliminary analysis in this direction showed that some trophic groups are more prone to extinction (Supplementary Information).

In conclusion, we demonstrated that the disruption of a keystone interaction by invasive species triggers changes to the structure of a multilayer mutualistic network, affecting multitrophic extinction cascades. While the loss of the keystone interaction erodes the community structure, it also sheds light on the importance of keystone interactions to restore communities. Management practices such as rewilding consider the reintroduction of extinct keystone species to their historical distribution to restore ecological functions [56]. Our findings highlight that such approaches could further benefit from considering keystone interactions and the multitrophic nature of ecological communities.

## Methods

### Study Area

The study was conducted in the Llao Llao Municipal Reserve (41^*◦*^9^*′*^*S*, 71^*◦*^18^*′*^*W*) and Nahuel Huapi National Park in Patagonia, Argentina (40^*◦*^58^*′*^*S*, 71^*◦*^31^*′*^*W*), located at the Subantarctic biogeographical region (Supplementary Information) [57]. Currently, 56% of the area of Nahuel Huapi National Park is occupied by the non-native ungulates: red deer (*Cervus elaphus*), dama deer (*Dama dama*), and domestic cattle (*Bos taurus*), which are the most abundant ungulates in the forest [58].

We leveraged this ongoing invasion scenario to compare non-invaded and invaded sites. We selected four 1-ha sites representing native plant communities, separated by more than 2 km (Supplementary Information). Two “intact” sites with the presence of the keystone interaction and no records of herbivory from ungulates. The two “invaded” sites were characterized by the presence of herbivory by non-native ungulates for over 100 years and historical records of the keystone interaction [45, 59]. Invaded sites have previous and current records of adult mistletoes and previous records of *D. gliroides*. The keystone interaction is ecologically extinct at invaded sites due to (i) the absence of mistletoe recruitment triggered by the herbivory of its main host (*A. chilensis*) and; (ii) changes to the vegetation, which produced the local extinction of its only seed disperser *D. gliroides* [18] (Supplementary Information).

### Data collection

We used data from Vitali et al. [19]. Briefly, at each site, we recorded pollinator-plant and frugivore-plant interactions for all flowering and fleshy-fruited plant species during two flowering and fruiting seasons (2017-2018 and 2018-2019; see Supplementary Information for sampling efforts). We recorded pollinator-plant interactions by conducting 10 minute censuses per plant. We considered an interaction when the visitor touched the reproductive structure of the flower. We recorded seed dispersal by birds by conducting one hour censuses per plant. We considered an interaction when the bird swallowed the fruit. To record seed dispersal by the marsupial *D. gliroides*, we captured individuals using Tomahawk traps, collected feces, and identified the consumed seeds. In addition, we accumulated more observation hours per plant species to improve sampling completeness of interactions by using infrared camera traps (Bushnell trophy cam).

We controlled for sampling effects by standardizing the sampling effort per species [19]. To this end, we conducted the same number of censuses of each plant species among sites (when the same species were present). Similarly, we standardized the sampling effort for the marsupial seed disperser and the use of camera traps among sites by trapping the same number of days and filming each plant species for the same amount of time (Tables S4,S5). The accumulation curves of species interaction richness indicate a high and consistent sampling completeness (Figs. S5,S6).

### Data analysis

Our goal was to compare intact and invaded sites. Therefore, we pooled interaction data across sites and seasons within each site type to increase our confidence in capturing the maximum number of interactions.

#### Connectivity between pollination and seed dispersal

We conducted two analyses to estimate the connectivity between pollination and seed dispersal. First, we recorded the presence of plant species *j* connecting both mutualisms and estimated its interlayer connectivity. We used two generalized linear mixed models (GLMMs) to test whether plant species’ presence and interlayer connectivity differed between the intact and invaded network. Our response variables were the presence/absence of a plant connecting layers and its interlayer link weight (*w*^*αβ*^). The explanatory variable was network types (intact vs invaded). We included “Species” as a random factor to control for species-specific differences. We used binomial and gamma with an inverse link function distribution because the response variables were binary and have positive continuous values not normally distributed, respectively [60]. We performed the analyses with the lme4 and glmmTMB packages in R software [61–63]. Second, we assessed if non-native ungulates change the connection between pollinator and seed disperser species. To do this, we built a binary matrix for the intact and invaded networks, encoding the existence of at least one indirect link between both trophic groups and calculated the proportion of indirect links per pollinator and seed disperser species (Fig. S3). Then, we performed a GLMM for each trophic group with the proportion of indirect links per species as a response variable and network types as an explanatory variable. We used “Species” as a random factor and gamma with an inverse link function distribution for the response variable.

#### Network modularity

We analyzed modularity using Infomap. Infomap detects the optimal network partition based on the movement of a random walker on the network and is specifically designed for multilayer networks [**?**, 64] (Supplementary Information). To assess if the observed network structure is a result of random processes we compared the observed number of modules to values generated by shuffling each network with a null model. To shuffle the networks, we used the “r2dtable” algorithm (vegan package within R [65]), modified to account for multilayer network nature. The algorithm shuffles the individual interactions within each layer, while preserving the total number of interactions per species (row and column marginal sums). Then, we calculated a p-value based on the proportion of the shuffled values that are larger or smaller than the observed value using a two-tailed t-test. Significant results indicate that the observed structure of each type of network is not random.

#### Species roles

We used the modularity results to assess the role that species play in network fragmentation and estimated the extent to which the invasion of non-native ungulates changed the role of species. To do this, we assigned species roles by calculating their “position” with respect to other species in their own module (within – module degree, “*z*”) and with species in other modules (among – module connectivity, “*c*”) [13] (Supplementary Information). According to *z* and *c* values, each species in each layer was assigned a role: peripheral (i.e., species with few links and mostly within the same module), connector (i.e., species with few links but linking several modules), module hub (i.e., species with several links within the same module), and network hub (i.e., species with several links within the same module and linking several modules). Therefore, plants which were present in both layers had two structural roles. In addition, for species that occurred in the intact and invaded networks, we recorded changes in structural role, for example a pollinator that was a connector in the intact network but peripheral in the invaded network.

#### Disturbance propagation via single species removal

In each simulation, we considered that the community reached an equilibrium when coextinctions did not propagate any longer after the removal of the target species [35]. Once equilibrium was reached, we calculated the total percentage of extinct species (*E*). Higher values of *E* indicate higher disturbance propagation. As the model included stochasticity, we repeated 100 times the removal of each species for each network and used the average value of *E*. We used a GLMM to test whether *E* was affected by the intact and invaded network, the structural role of the removed species (peripheral, connector, module hub, and network hub) or the interaction between them. We included “Species” as a random factor and assumed gamma distribution in the model. In addition, to compare among different levels of the explanatory variables, we performed a multiple comparison test. Analyses were performed using R software and lme4, multcomp and lsmeans packages [66]. See details in Supplementary Information.

#### Network robustness via sequential removal

For each network, we removed species sequentially under three scenarios, according to their structural role, which is an estimate for the species’ ability to maintain network cohesiveness: (i) removal from the most to the least connected structural role (order: network hub, module hub, connector, peripheral); (ii) the opposite order; and (iii) random removal order. We used the stochastic extinction model explained above but this time, in each scenario, we removed the first species and once the community reached equilibrium, we removed the next species, and so on until all species of the network became extinct. We calculated the robustness of the network to species extinction as the area under the curve (AUC) of the species lost against species present in the network. Higher values of AUC suggest higher tolerance to sequential loss of species [67]. For each network and scenario combination, we ran 1000 simulations and calculated the AUC. Then, we used a GLM model to test if AUC (response variable assuming gamma distribution) was affected by the intact and invaded network, removal scenario and the interaction between them. In addition, we performed a multiple comparison test to compare among different levels of the explanatory variables. Analyses were performed using the lme4 and lsmeans packages in R. Finally, we controlled for network size because it is correlated with other structural properties [44] and could therefore affect *E* and AUC. We did that by bootstrapping the largest network. Network size affected *E* but not AUC (Supplementary Information).

## Supplementary information

Supplementary Figures, Tables and Methods.

## Acknowledgments

We thank the staff of Nahuel Huapi National Park, D. Mujica and C. Chehebar, the staff of Parque Municipal Llao-Llao and Dirección de Áreas Protegidas, M.S. Millerón for logistic support and permission to carry out fieldwork. We also thank A.P. Duarte, A. Fernández, A. Santone, B.R. Delgado, B. Lovazzano, E. Valfosca, I. Villa, J.G. Calzada, J. Gastaudo, K. Buteler, L.M. Valfosca, M.E. Valfosca and P. San Pedro for valuable assistance in the field. We are thankful to J.G. Calzada for creating the illustrations in Fig 1. This research was supported by grants from “Agencia Nacional de Promoción Científica y Tecnológica” of Argentina (PICT 2014-2484) to MARC, “The Rufford Foundation” to AV (ID 26510-1), and the “Israeli Science Foundation” (number 1281/20) to SP. This work also was supported by a scholarship from the Israeli Council of Higher Education to AV.

## Data and code availability

Raw data supporting the results and code will be available online upon acceptance.

## Notes

### Competing Interest Statement

The authors have declared no competing interest.

