## Supplementary Information for "Invasive species modulate the structure and stability of a multilayer mutualistic network"

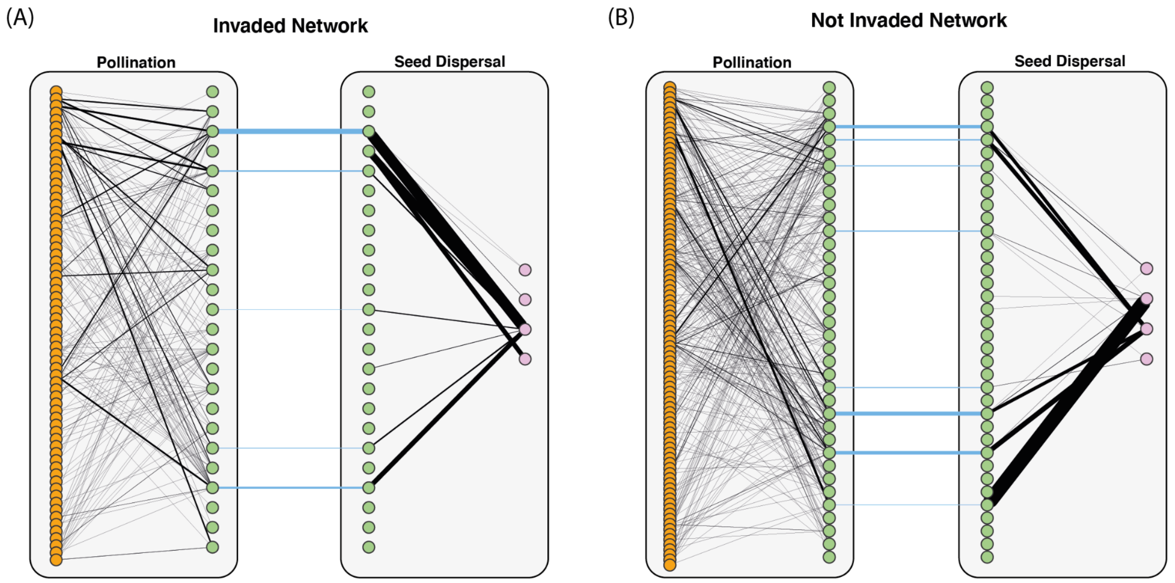


**Figure S1**. The structure of the pollinator-plant-seed disperser multilayer networks. The networks were constructed using the data gathered from (A) invaded plots, and (B) non-invaded ( intact) plots. Each network is composed of two layers, represented by rectangles. The layer on the left represents pollination and on the right, seed dispersal. Each layer contains intralayer links (black lines) representing interactions between pollinator (orange) and plant (green) species or between plant (green) and seed disperser (pink) species. The weight of the intralayer edges (range 0-1) is the relative number of visits of pollinators or seed dispersers to plants out of all the links in the respective layer. Interlayer edges connect the same plant to itself in both layers. Interlayer link weight (range 0-1) is the total number of indirect links between seed dispersers and pollinators mediated by the plant, out of all indirect links. Plants that do not interact in both layers are drawn as singletons but do not influence calculations.

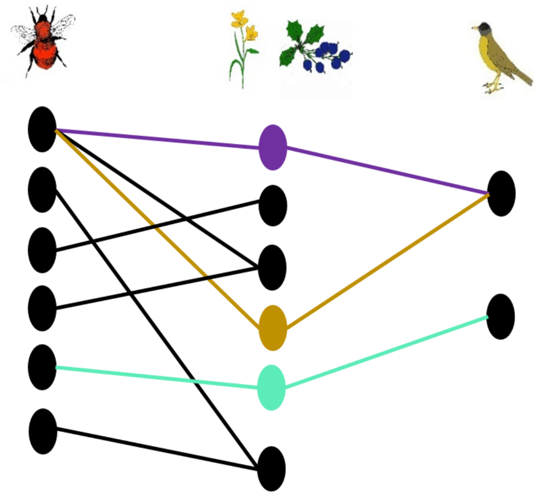

**Figure S2**. Example of indirect links between pollination and seed dispersal mutualism. Circles from left to right represent pollinator, plant, and seed disperser species. Three indirect links are represented by violet, brown and light blue lines. Each indirect link arises when a plant is pollinated and dispersed by animals. Two indirect links of a plant species will be considered different when at least the pollinator or seed disperser species of the indirect link differs. For example, if a plant is pollinated by three species and dispersed by two species, the plant will have six different indirect links.

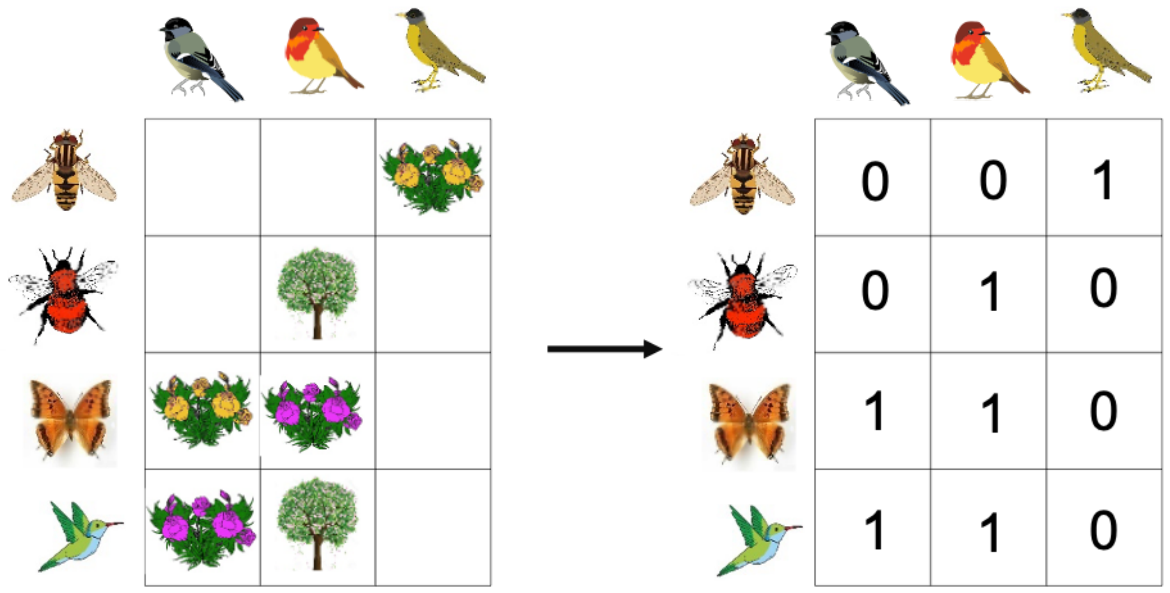


**Figure S3**. Matrix of indirect links between pollinators and seed dispersers. To build the matrix for the intact and invaded networks, we first detected whether a plant species was visited by a pollinator and a seed disperser species. Then, we assigned the presence of an indirect link between a pollinator (row) and a seed disperser (column) when they visited the same plant species (“1”). On the contrary, we assigned the absence of an indirect link (“0”) for those pollinator and seed disperser species that were not detected visiting the same plant species. We used a binary matrix because few pollinator and seed disperser species shared more than one plant species. Then, we calculated the proportion of indirect links per pollinator species as row marginal sum and row marginal sum divided by the total number of seed dispersers, respectively. For seed disperser species, we calculated the proportion of indirect links in the analogous way.

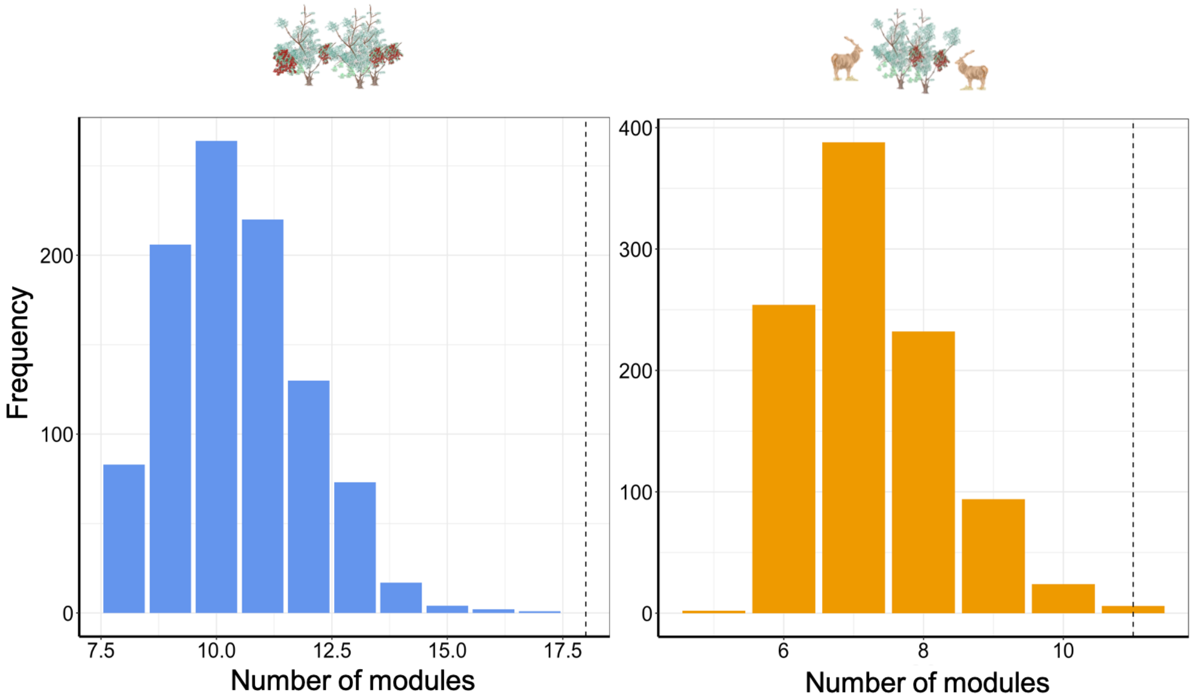


**Figure S4**. Comparison of the observed number of modules to values generated by shuffling the network for both types of networks. Vertical dotted line indicates the observed number of modules in the empirical network. Left and right panels indicate the intact and invaded site, respectively.

### Supplementary Methods: Site description and sampling effort

#### Description of sites:

To characterize each site, we delimited three transects of 100m separated from each other by 25m. In each transect, we evaluated the vegetation structure, understory cover, plant richness and abundance of *Aristotelia chilensis* and *Tristerix corymbosus* by establishing five circular plots of 2m radius. Vegetation structure was estimated by counting the number of branches (5-10 cm in diameter and ≤ 45° with respect to the ground) that touched a 3m vertical pole located in the center of the circular plots. Understory cover was estimated by measuring four times with a densitometer placed in the center of the circular plots. In addition, we also established five transects of 100m x 2m in each site and counted the number of *T. corymbosus* flowers during its flowering period. At site level, the values of plant richness, abundance of *A. chilensis* and *T. corymbosus,* and number of *T. corymbosus* flowers were accumulated considering all the plots and transects. On the other hand, the values of vegetation structure and understory cover per site were calculated by averaging the values of the plots (Table S1).

**Table S1.** Description of sites.

| Type of site | Site | Plant richness | Vegetation structure[1] | Understory cover[1] | *A. chilensis* abundance per ha | *T. corymbosus* abundance per ha | Number of *T. corymbosus* flowers per ha |
|---|---|---|---|---|---|---|---|
| NI | 1 | 21 | 2.60±0.19 | 0.89±0.01 | 39 | 29 | 8830 |
| | 2 | 22 | 2.60±0.23 | 0.88±0.01 | 36 | 22 | 7220 |
| I | 1 | 13 | 0.73±0.15 | 0.82±0.03 | 21 | 4 | 967 |
| | 2 | 16 | 0.60±0.16 | 0.89±0.01 | 19 | 3 | 1360 |

Notes: NI: intact sites; I: sites invaded by non-native ungulates. [1]Values represent means ± standard error. Notes: Table modified from [1].

To measure the presence of non-native ungulates, we monitored with camera traps each study site. We placed camera traps 1m above the ground to record 30 seconds during day and night. Cameras were triggered by movement. A record of non-native ungulates was considered when we recognized its identity from the videos. We estimated the proportion of camera trap videos that recorded ungulates at each site as a quantitative measure of herbivore occupancy. In addition, we randomly selected 50 individuals of *A. chilensis* which we scanned for evidence of damage (i.e., presence or absence) to the leaves and branches. From the same individuals, we counted the total number of seedlings of *T. corymbosus* to estimate the effect of non-native ungulates on its recruitment (Table S2).

**Table S2.** Herbivory by non-native ungulates and its effect on mistletoe recruitment in each site.

| Type of site | Site | Percentage of videos that recorded non-native ungulates | Percentage of damaged *A. chilensis* per 50 individuals | Number of *T. corymbosus* seedlings in 50 individuals of *A. chilensis* |
|---|---|---|---|---|
| NI | 1 | 0 | 0 | 42 |
| | 2 | 0 | 0 | 49 |
| I | 1 | 54.5 | 98 | 0 |
| | 2 | 50 | 82 | 0 |

Notes: NI: intact sites; I: sites invaded by non-native ungulates. Table modified from [1].

**Sampling effort:**

**Table S3.** Sampling efforts of pollination network of each site.

| Type of site | Year | Site | Total censuses | Time (hs) | No. plant species | No. visitors species | No. links | No. interactions |
|---|---|---|---|---|---|---|---|---|
| NI | 2017 2018 | 1 | 486 | 81 | 26 | 44 | 110 | 485 |
| | | 2 | 361 | 60.2 | 18 | 39 | 93 | 622 |
| | 2018 2019 | 1 | 442 | 73.7 | 24 | 54 | 138 | 1046 |
| | | 2 | 306 | 51 | 16 | 47 | 88 | 681 |
| I | 2017 2018 | 1 | 211 | 35.2 | 11 | 35 | 63 | 486 |
| | | 2 | 324 | 54 | 12 | 24 | 36 | 200 |
| | 2018 2019 | 1 | 185 | 30.8 | 10 | 27 | 53 | 411 |
| | | 2 | 333 | 55.5 | 13 | 29 | 50 | 420 |

Notes: NI: intact sites; I: sites invaded by non-native ungulates. Table modified from [1].

**Table S4.** Sampling efforts of seed dispersal network of each site

| Type of site | Year | Site | Total censuses | Time (hs) | Camera traps (hs) | Tomahawk traps (days) | No. plant species | No. visitors species | No. links | No. interactions |
|---|---|---|---|---|---|---|---|---|---|---|
| NI | 2017 2018 | 1 | 25 | 25 | 1440 | 8 | 6 | 2 | 8 | 294 |
| | | 2 | 39 | 39 | 1440 | 8 | 7 | 3 | 10 | 299 |
| | 2018 2019 | 1 | 36 | 36 | 1944 | 8 | 8 | 3 | 10 | 374 |
| | | 2 | 28 | 28 | 1680 | 8 | 7 | 3 | 12 | 291 |
| I | 2017 2018 | 1 | 27 | 27 | 1200 | 8 | 4 | 2 | 5 | 201 |
| | | 2 | 15 | 15 | 1200 | 8 | 2 | 2 | 3 | 71 |
| | 2018 2019 | 1 | 39 | 39 | 1200 | 8 | 6 | 4 | 9 | 213 |
| | | 2 | 23 | 23 | 1200 | 8 | 4 | 1 | 4 | 191 |

Notes: NI: intact sites; I: sites invaded by non-native ungulates. Table modified from [1].

**Table S5.** Sampling effort of camera traps per plant species in each site.

| **2017-2018** | | | | | | | | | |
|---|---|---|---|---|---|---|---|---|---|
| Type of site | Site | Camera traps (hours) per species | | | | | | | |
| | | *Berberis darwinii* | *Schinus patagonicus* | *Tristerix corymbosus* | *Prunus avium* | *Aristotelia chilensis* | *Relbunium hypocarpium* | *Ribes magellanicum* | *Azara microphylla* |
| NI | 1 | 240 | 240 | 240 | 240 | 240 | 240 | - | - |
| | 2 | 240 | - | 240 | - | 240 | 240 | 240 | 240 |
| I | 1 | 240 | 240 | 240 | - | 240 | - | - | 240 |
| | 2 | 240 | 240 | 240 | - | 240 | 240 | - | - |
| **2018-2019** | | | | | | | | | |
| Type of site | Site | Camera traps (hours) per species | | | | | | | |
| | | *Berberis darwinii* | *Schinus patagonicus* | *Tristerix corymbosus* | *Prunus avium* | *Aristotelia chilensis* | *Relbunium hypocarpium* | *Ribes magellanicum* | *Azara microphylla* |
| NI | 1 | 240 | 240 | 240 | 264 | 240 | 240 | 240 | 240 |
| | 2 | 240 | 240 | 240 | - | 240 | 240 | 240 | 240 |
| I | 1 | 240 | 240 | 240 | - | 240 | - | - | 240 |
| | 2 | 240 | 240 | 240 | - | 240 | 240 | - | - |

Notes: NI: intact sites; I: sites invaded by non-native ungulates. Table modified from [1].

**Table S6.** Overall network properties of each type of site.

| Type of site | Layer | Plant richness | Pollinator/Seed dispersal richness | Connectance[2] |
|---|---|---|---|---|
| NI | Pollination | 37 (8)[1] | 95 | 0.093 |
| | Seed dispersal | | 4 | 0.45 |
| I | Pollination | 24 (5)[1] | 67 | 0.113 |
| | Seed dispersal | | 4 | 0.393 |

Notes: NI: intact site; I: site invaded by non-native ungulates. [1]Values indicate the number of plant species present in both layers. [2]Calculated as the sum of links divided by the number of cells in the matrix.

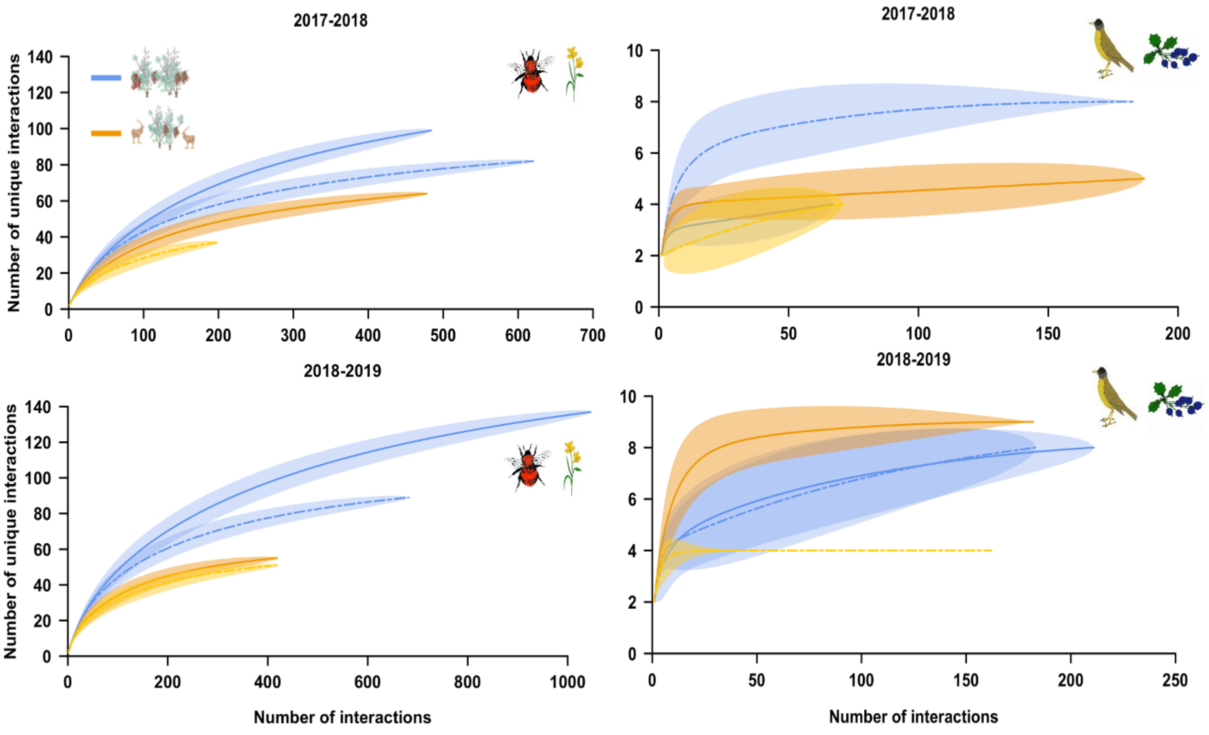


**Figure S5.** Smoothed accumulation curves of species interaction richness according to number of recorded interactions in each site and season. Left and right panels represent interactions of pollinators visiting flowers and birds removing fruits, respectively. Line color represents site type: intact and invaded in light blue and orange, respectively. Continuous and dashed lines represent different sites. Each line shows the average of species interaction richness represented by the number of recorded interactions. Shaded area around each curve indicates the 95% confidence interval around the mean. The smoothing process applied to accumulation curves is optimal to reduce the stochastic noise produced by temporal and spatial biases on sampling effort distribution and to avoid specific bias produced by the order in which censuses were performed [2]. Figure modified from [1].

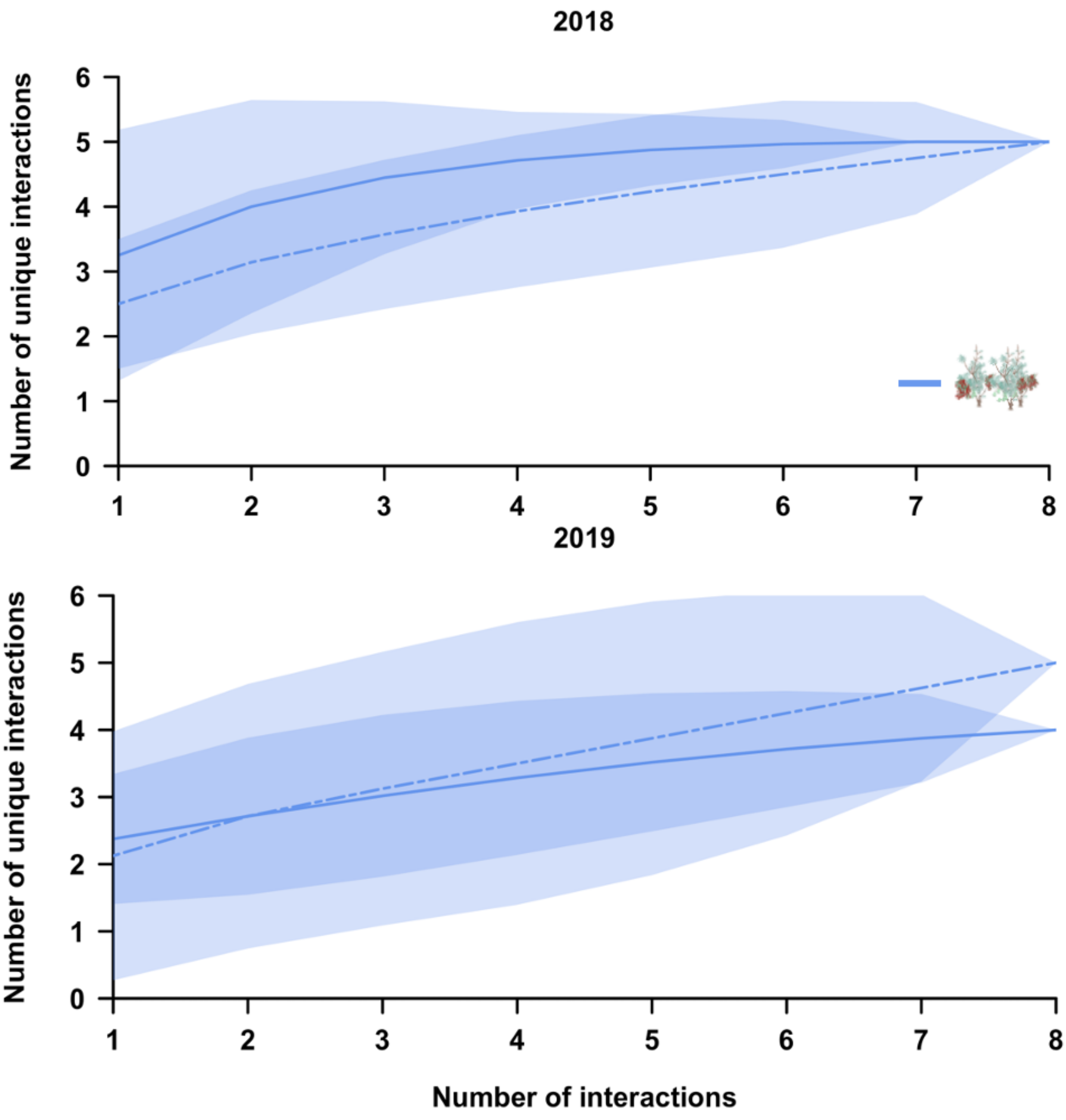


**Figure S6.** Smoothed accumulation curves of interactions richness recorded during sampling of the seed disperser marsupial (*Dromiciops gliroides*) according to the number of sampling days for each intact site and season. Continuous and dashed lines represent different sites. Each line shows the average of species interaction richness represented by the number of sampled days. Shaded area around each curve indicates the 95% confidence interval around the mean. Invaded sites were excluded from the figure because we did not capture individuals of *D. gliroides*. Figure modified from [1].

### Supplementary Methods: Additional explanations of data analysis

#### Modularity:

We identified groups of tightly connected species with a modularity analysis using Infomap [3,4]. Infomap detects the optimal network partition based on the movement of a random walker on the network and is specifically designed for multilayer networks (see [3,5,6] for details). For any given partition of the network, the random walker moves across nodes in proportion to the weight of the edges. The amount of information it costs to describe the walk is quantified using the objective function $L$ called the map equation. The optimal network partition is the one that minimizes $L$. In multilayer networks, nodes representing observable entities such as plants are called physical nodes, and nodes describing the occurrence of species in the layers are called state nodes. The random walker moves from state node to state node within and across the layers on intra- and inter-links respectively. Therefore, the multilayer network representation is not merely an extended network with unique nodes in all layers and a module can encompass plants, pollinators, and seed dispersers.

#### Structural role of species:

We classified the species according to their structural role following [7]. The role of a species was defined by its standardized within-module degree z (i.e., how well the species is connected with other species in the same module) and its among-module connectivity c (i.e., how well the species connects species from different modules):

$$z = \frac{k_{is} - k_s}{SD_{ks}}$$

$$c = 1 - \sum_{t=1}^{N_M} \left(\frac{k_{it}}{k_i}\right)^2$$

In z, $k_{is}$ is the number of links of $i$ to other species in its own module $s$, $k_s$ is the average of within-module degree of all species in $s$, and $SD_{ks}$ is the standard deviation of within-module degree of all species in s. In c, $k_{it}$ is the number of links from $i$ to species in the module t, $k_i$ is the degree of the species $i$, and $N_M$ is the total number of modules. Values of c can range between 0 and 1, being 0 when all the links of the species $i$ are within its own module and 1 when its links are distributed evenly among modules.

Using z and c values, we sorted all the species into different structural roles: peripheral, connector, module hub, or network hub. A peripheral species has few links and mostly within the same module (a low $z \leq 2.5$ and a low $c \leq 0.62$, light blue circles in Fig. S1). A connector species has few links but links several modules (a low $z \leq 2.5$ and a high $c > 0.62$, pink circles in Fig. S1), being important for the network cohesion because it glues modules together. A module hub has several links within the same module and is important for the integrity of its own module (a high $z > 2.5$ and a low $c \leq 0.62$, purple circles in Fig. S1). A network hub has several links within the same module and between species of different modules (a high $z < 2.5$ and a high $c > 0.62$, orange circle in Fig. S1), being important for the integrity of the whole network. Classification of all species according to their structural role in the Fig. S2.

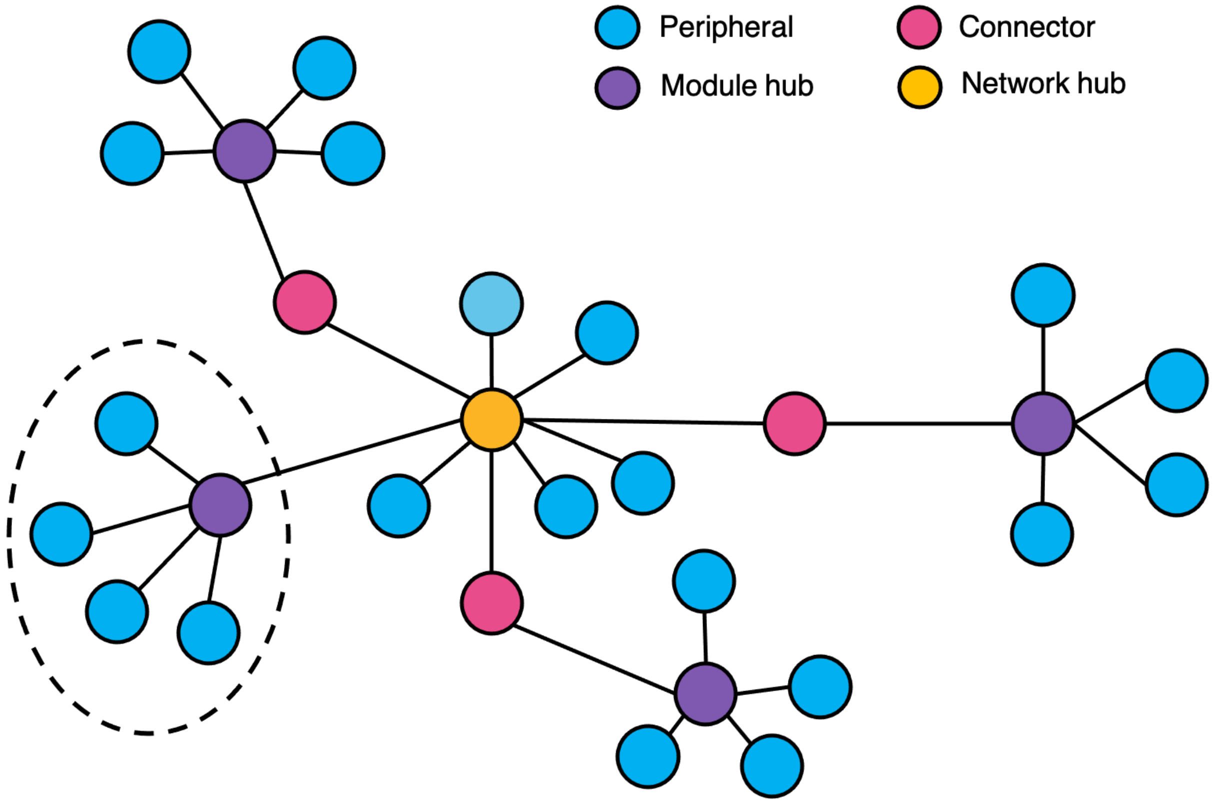


**Figure S7.** Example of species classification according to their structural role. Circles and lines represent species and links between species, respectively. Color of circles indicates the structural role of species: peripheral (light blue), connector (pink), module hub (purple), network hub (orange). Dotted line delimits a module.

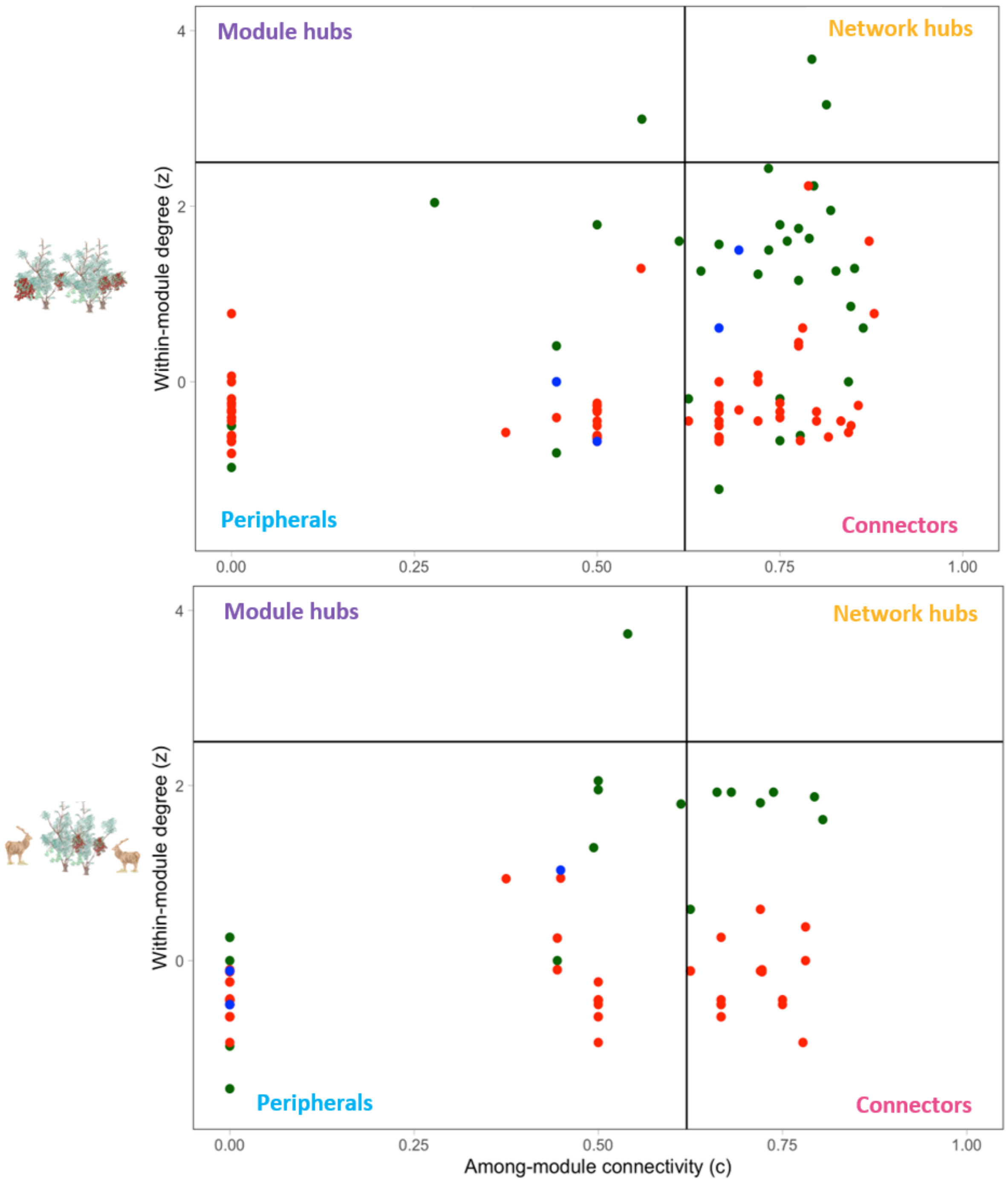


**Figure S8.** Distribution of species according to their structural role in the intact (upper panel) and invaded (lower panel) network. Each point represents a species and its color indicates its trophic group: pollinator (red), plant (green), and seed disperser (blue).

**Stochastic coextinction models:**

We modified the "netcascade" function, elaborated by [8], to include different trophic groups in the stochastic coextinction models (SCM). In our SCM models, a simulation begins with a primary extinction, which can lead to coextinction events of species. The probability of species $y$ to suffer coextinction after the removal of its mutualistic partner $x$ is $P_{yx} = d_{yx} * R_y$. $d_{yx}$ is the dependence of species $y$ on its partner $x$ and is defined as the number of visits recorded between $y$ and $x$ divided by the total number of visits involving species $y$ with species of the same trophic group as $x$ in the network [9]. Thus, the formula of $d_{yx}$ will change according to the trophic groups of both species involved in the extinction chain. In addition, $R_y$ is a constant, which indicates the intrinsic demographic dependence of species $y$ on the mutualism to which $x$ belongs. Because plant species may interact with both pollinator and seed disperser species, thus, they have two values of R according to their demographic dependence on pollination and seed dispersal by animals, respectively. When a coextinction event occurs, each remaining species in the other trophic group has a probability to suffer extinction (if $P_{yx} > 0$). In each step, the dependences between species $y$ and $x$ ($d_{yx}$) are recalculated.

For each extinction scenario, the probability of species $y$ to suffer coextinction after the removal of its mutualistic partner $x$ will be:

- If $x$ is a pollinator species, the probability of co-extinction of a plant $y$ is: $P_{yx} = d_{yx} * R_{y\alpha}$ and $d_{yx} = \frac{W_{yx}}{\Sigma W_{y\alpha}}$, where $R_{y\alpha}$ is how much the plant $y$ depends on pollination by animals, $W_{yx}$ is the number of visits between $y$ and $x$, and $\sum W_{y\alpha}$ is the total number of visits of $y$ with pollinators.

- If $x$ is a seed disperser species, the probability of co-extinction of a plant $y$ is: $P_{yx} = d_{yx} * R_{y\beta}$ and $d_{yx} = \frac{W_{yx}}{\Sigma W_{y\beta}}$, where $R_{y\beta}$ is how much the plant $y$ depends on seed dispersal by animals, $W_{yx}$ is the number of visits between $y$ and $x$, and $\sum W_{y\beta}$ is the total number of visits of $y$ with seed disperser species.

- If $x$ is a plant species, the probability of co-extinction of a pollinator or seed disperser species $y$ is: $P_{yx} = d_{yx} * R_y$ and $d_{yx} = \frac{W_{yx}}{\Sigma W_y}$, where $R_y$ is how much the pollinator or seed disperser $y$ depends on resources provided by plants (e.g., nectar for pollinators and fruits for seed dispersers), $W_{yx}$ is the number of visits between $y$ and $x$, and $\sum W_y$ is the total number of visits of $y$ with plant species.

**Estimation of the intrinsic demographic dependence of each species (R):**

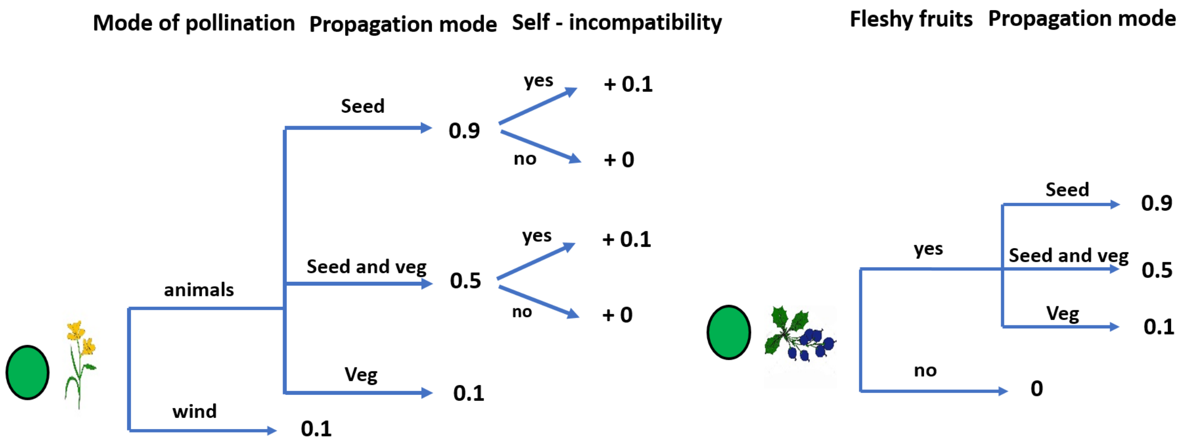


**Figure S9.** Diagram to classify the R of plant species according to its dependence on pollination (left panel) and seed dispersal mutualisms (right panel). For each plant species, we scored the dependence on pollination mutualism according to its mode of pollination (e.g., by wind or animal), its propagations mode (seed, seed and vegetative, or vegetative) and if the plant had a self-incompatibility breeding system. For example, a plant species with a self-incompatibility breeding system, pollinated by animals and which propagates only by seeds will have a strong dependence on pollination mutualism (R = 1). Plants pollinated by wind were scored as 0.1 because they are occasionally visited by insect pollinators [10]. In addition, we scored the dependence on seed dispersal mutualism of each plant species according to its production of fleshy fruits (i.e., “yes” or “not”) and its propagation mode. For example, plant species producing fleshy fruits, which propagate only by seeds will have a strong dependence on seed dispersal mutualism (R = 1).

Following [11], for each flower visitor species, we scored its dependence on plants as "high" (R = 0.9) to flower visitors with pollen and nectar as obligate resource in its diet, as "medium" (R = 0.5) to those species likely consuming alternative food resources (e.g., insects, dung), and as "low" (R = 0.1) to those species which were observed visiting flowers in our study but rarely consume nectar or pollen (Fig. S4). In the cases that we could not identify the flower visitor to species level or no information was available, we used information of the closest taxonomic level (e.g., genera, family). When we could identify the flower visitor to order level at most, we randomly scored the species as "medium" or "low". For each seed disperser species, we scored its dependence on plants as "high", "medium" or "low" (R = 0.9, 0.5, or 0.1, respectively) according to its diet (Fig. S4). We identified the dietary components of each species (e.g., fruits, seeds, invertebrates, etc.) and ranked them according to their relevance to the species diet following [12]. For example, we scored as high dependence on plants to those animal species that mainly consume fleshy fruits and had less than three dietary components. Classification of species according to their dependence on mutualism and all information needed to classify them were obtained from a comprehensive literature review (Table S1-S4 and list of reviewed articles at the end of the appendix).

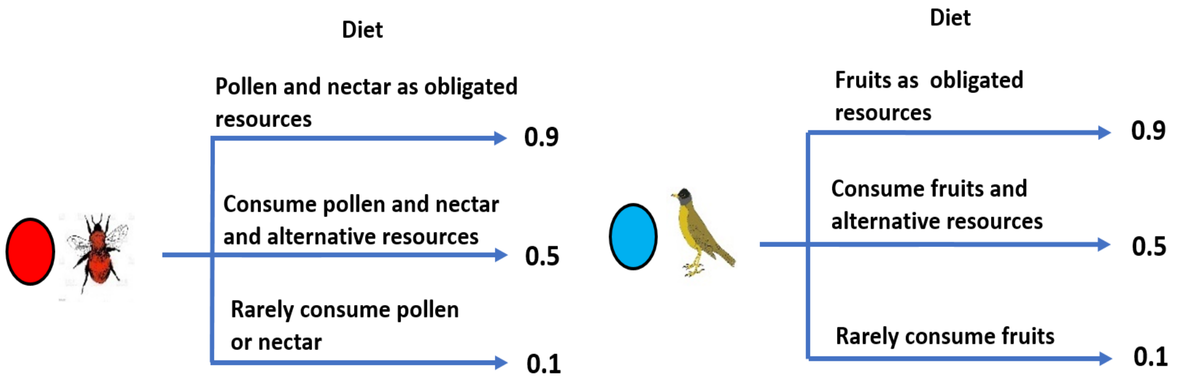


**Figure S10.** Classification of the R of pollinator and seed disperser species (left and right panels) according to dependence on plant resources. Full explanation in the main text above.

**Sensitivity of extinction simulations to R:**

We performed a sensitivity analysis to test if the patterns found in the extinction models were due to the dependence on the mutualism (R) assigned to species. To do this, we created scenarios where we reduced and increased the values of R of each species used in the simulations by 0.1 until all species had either no or full dependence on the mutualisms. In one extreme scenario, all species have no dependence on the mutualistic interactions and the removal of a species will not produce cascading effects. In the opposite extreme scenario, all species are completely dependent on mutualistic interactions (R = 1). This last scenario represents a typical extinction model that does not incorporate the dependence of species on the mutualisms because the probability of species extinction ($P_{yx} = d_{yx} * R_y$) will only depend on the interactions between them ($d_{yx}$). For each scenario, we ran the same simulations described in the main text to estimate the total percentage of extinct species and the robustness (AUC) after a single species removal and the sequential removal of species according to their structural role, respectively.

The outputs of the extinction models (total percentage of extinct species and AUC) were quantitatively sensitive to changes in R. However, qualitatively the patterns found in the results remained unchanged (Fig. S5 and S6). Specifically, we find that the effect of species roles on the percentage of extinct species and the AUC remains the same across all values of R (the lines in Figures S5 and S6 never cross each other).

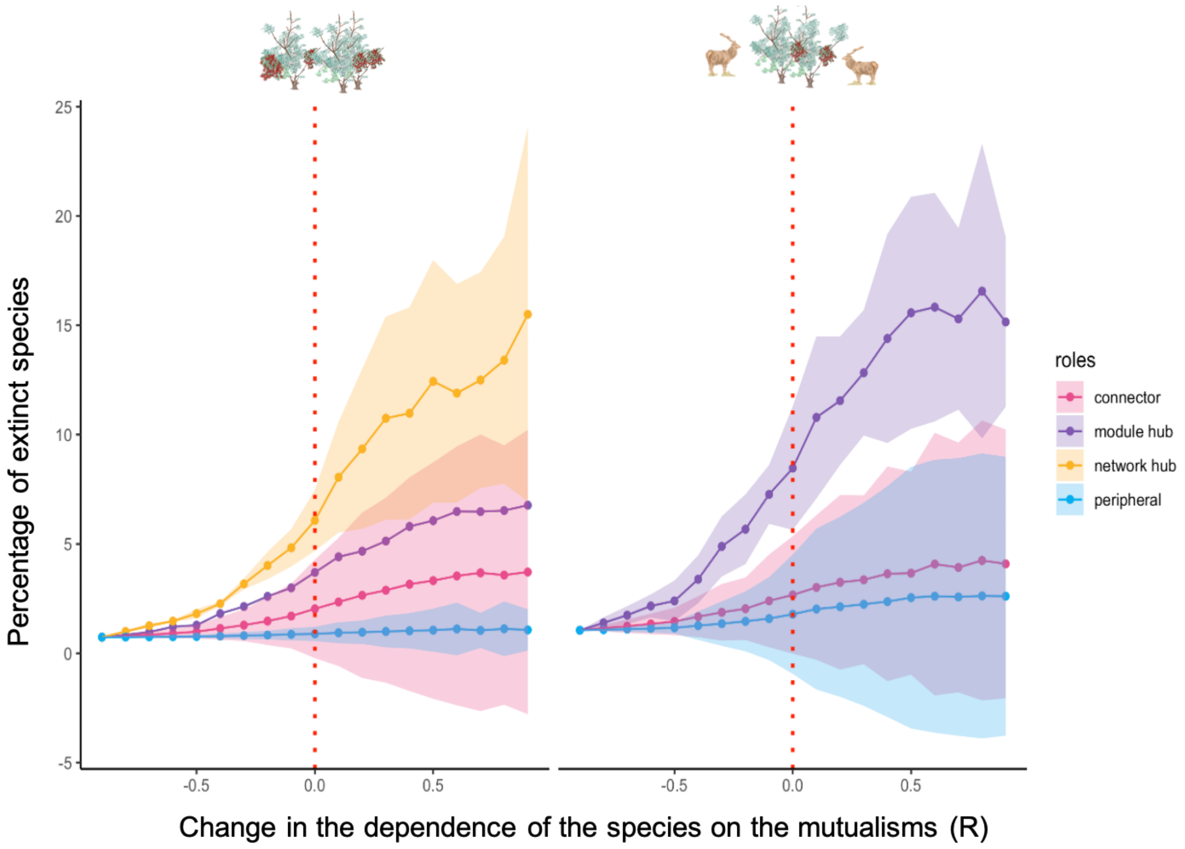


**Figure S11.** Percentage of extinct species according to changes in the dependence of the species on the mutualisms (R) and the structural role of the removed species. Values close to -1 and 1 on the horizontal axis indicate no and full dependence of all species on the mutualistic interactions, respectively. Each line indicates the percentage of extinct species after removing a network hub (orange), module hub (purple), connector (pink), and peripheral species (light blue). Shaded area around each line indicates the 95% confidence interval around the mean. Vertical red dotted line indicates the observed percentage of extinct species estimated by the assigned values of R to each species in the main text. Left and right panels represent the intact and invaded network respectively.

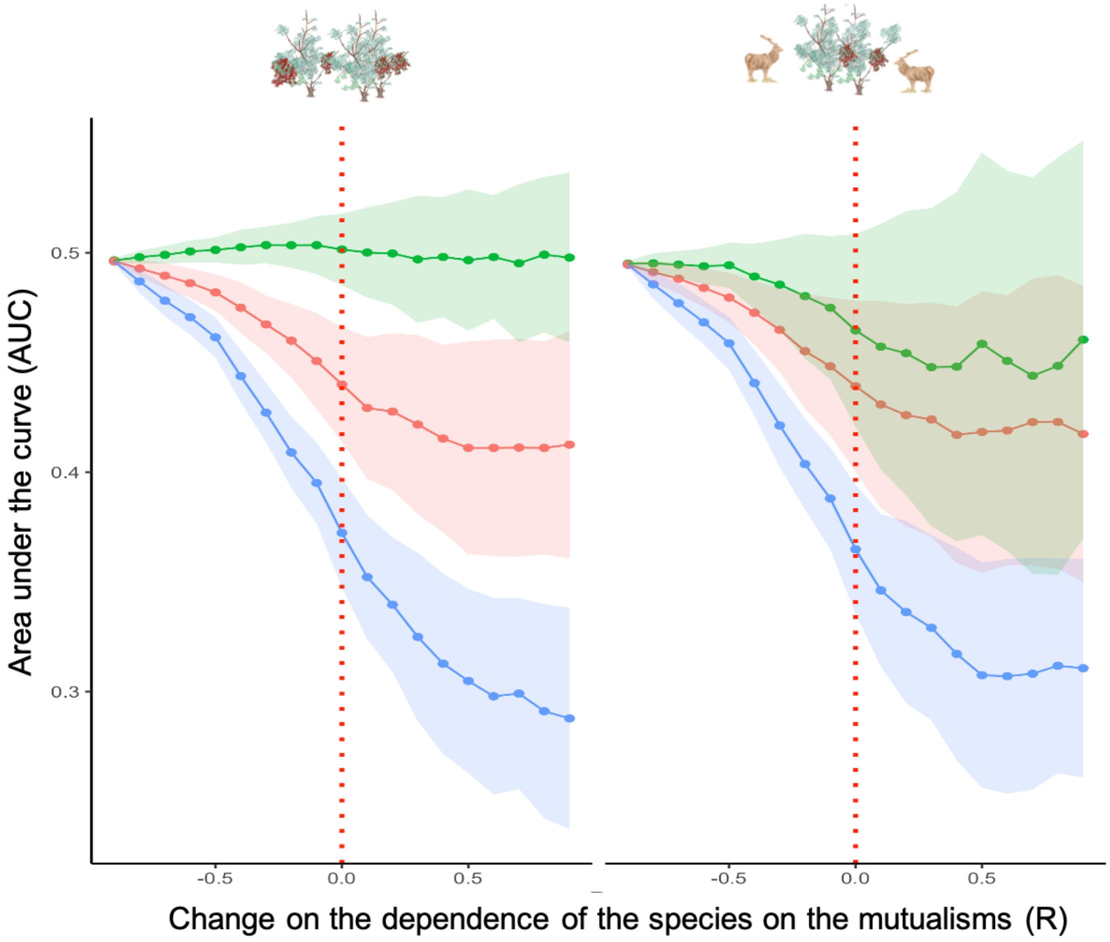


**Figure S12.** Area under the curve (AUC, proxy of robustness) according to changes in the dependence of the species on the mutualisms (R) and the order of species removal. Values close to -1 and 1 on the horizontal axis indicate no and full dependence of all species on the mutualistic interactions, respectively. Each line indicates robustness of the community after sequentially removing all species from the most to the least connected structural role (blue, order: network hub, module hub, connector, peripheral), from the least to the most connected structural role (green, order: peripheral, connector, module hub, network hub), and at random (light red). Shaded area around each line indicates the 95% confidence interval around the mean. Vertical red dotted line indicates the observed robustness estimated by the assigned values of R to each species in the study. Left and right panels represent the intact and invaded network.

**Extinction analysis controlling for network size:**

We controlled for network size because it is correlated with other network properties, such as modularity [13], and could therefore affect the outputs of the simulation model (total percentage of extinction and robustness). To do this, we first bootstrapped the largest network (intact network) 300 times to obtain in each trophic group the same number of species as in the counterpart trophic groups of the smallest network (invaded network). Second, for each simulated network, we calculated the total percentage of extinct species after a single species removal and the robustness (AUC) after sequentially removing species according to their structural role (see the main text for more details). Then, we tested if these variables changed between the intact network controlled for size and the intact network without controlling for size. We performed a GLMM with the total percentage of extinct species as response variable and treatment (controlled network vs. uncontrolled network) as fixed factor. We included "Species" as a random factor in the model. In addition, we performed a GLM with the area under the curve (AUC, proxy of robustness) as response variable and treatment as fixed factor. We assumed that response variables followed a gamma distribution in both models [14]. All the analyses were performed using R software and the "lme4" package.

We found that the total percentage of extinct species was affected by network size (F = 35.18, $P < 0.001$; Fig. S7) but not the robustness (F = 0.26, $P = 0.61$; Fig. S8). The total percentage of extinct species increased 1.4 times after controlling for network size, indicating a greater propagation of disturbances (Fig. S7).

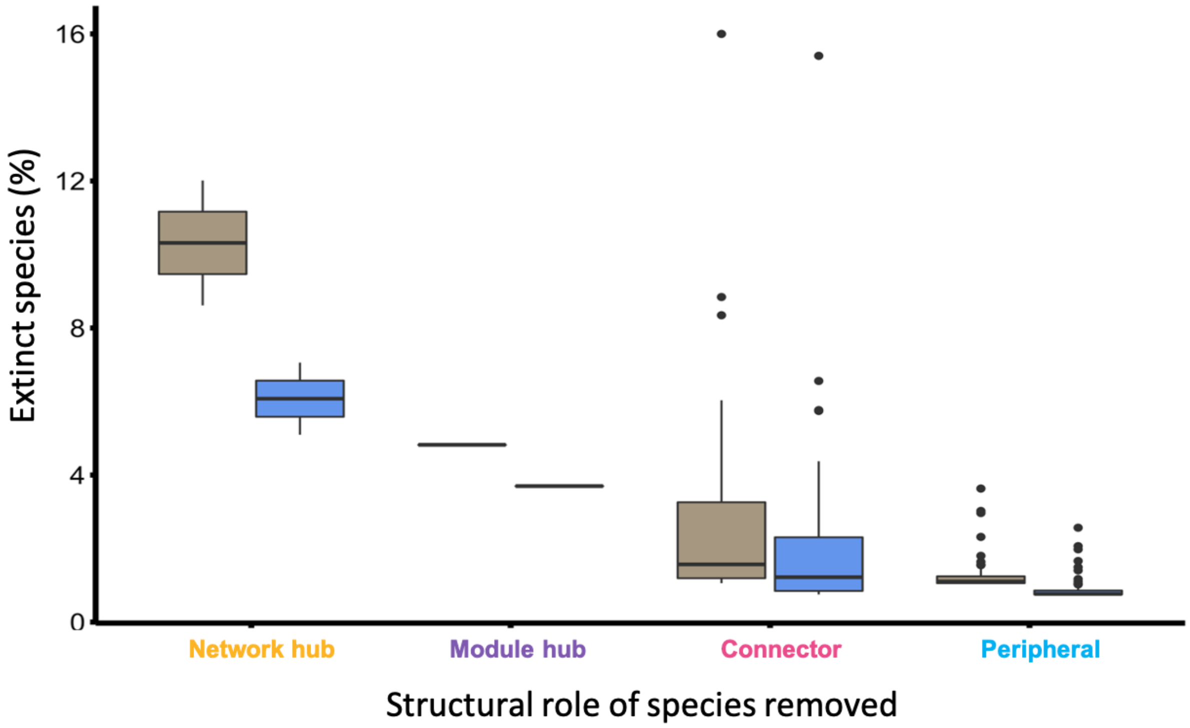


**Figure S13.** Percentage of extinct species in the intact network after the removal of a single species according to their structural role and after controlling for size. Color of box plots depicts: intact network controlled for size (gray) and intact network without controlling for size (light blue).

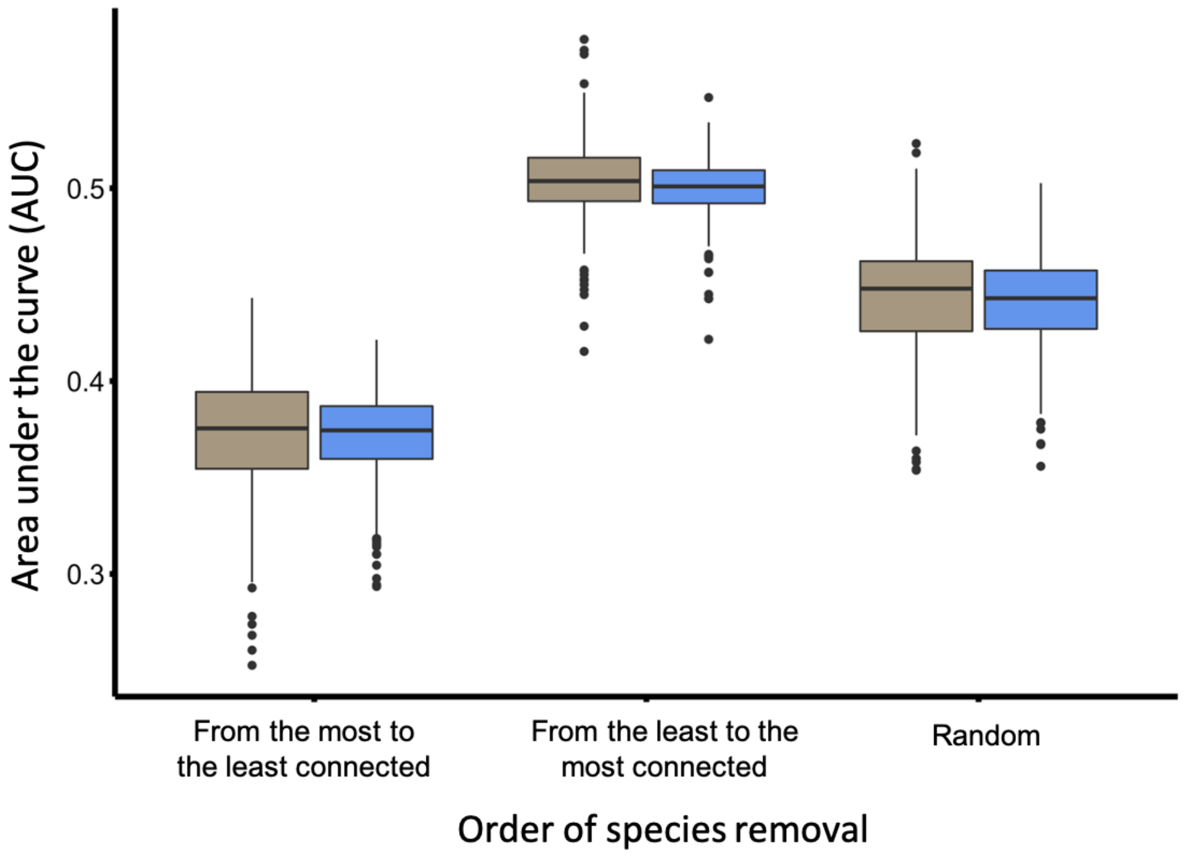


**Figure S14**. Area under the curve (AUC, proxy of robustness) in the intact network after sequentially removing species according to their structural role and controlling for size.

Color of box plots depicts: intact network controlled for size (gray) and intact network without controlling for size (light blue). Horizontal axis indicates the order of species removal: from the most to the least connected structural role (order: network hub, module hub, connector, peripheral), from the least to the most connected structural role (order: peripheral, connector, module hub, network hub), and at random.

**Disturbance propagation according to trophic groups:**

We calculated the total percentage of extinct species of each trophic group after simulating a single species removal in the stochastic extinction model (see main text for more details). To test whether certain trophic groups are more susceptible to the extinction of a particular trophic group in the intact and invaded network, we built three models using a combination of response variables (percentage of extinct pollinator, plant, or seed disperser species) and the interaction between invasion status (intact or invaded network) and removed trophic group (pollinator, plant, or seed disperser) as explanatory variables. At first, all models included interaction between explanatory variables, however, we discarded it when it was non-significant. We included "Species" as a random factor and assumed gamma distribution in the models [14]. In addition, when we detected a significant effect in explanatory variables with more than two levels and a significant interaction between explanatory variables, we performed a post hoc test using the False Discovery Rate (FDR) method in order to compare the response variables among all the levels [15] and a multiple comparison test. All the analyses were performed using R software and lme4, multcomp and lsmeans packages [16,17].

Only the percentage of pollinators that went extinct after the removal of a species differed statistically between the intact and invaded network, being almost 1.7 times higher in the invaded network (GLMM, $X^2_{1, 231} = 7.58$, $P < 0.01$; asterisk in Fig. S9). In addition, pollinator, plant, and seed disperser species were less susceptible to extinction when a pollinator species was removed. After the removal of a pollinator species, the percentage of pollinators and plants that went extinct were almost 2 and 6.6 times lower than after the removal of a plant (post-hoc test, z = 3.690, $P < 0.01$; z = 7.205, $P < 0$ .01) and 3.6 and 11.6 times lower than after the removal of a seed disperser species (post-hoc test, z = 2.544, $P <$ 0.027; z = 7.381, $P < 0.01$). Seed dispersers were not affected by the extinction of pollinators (Fig. S9).

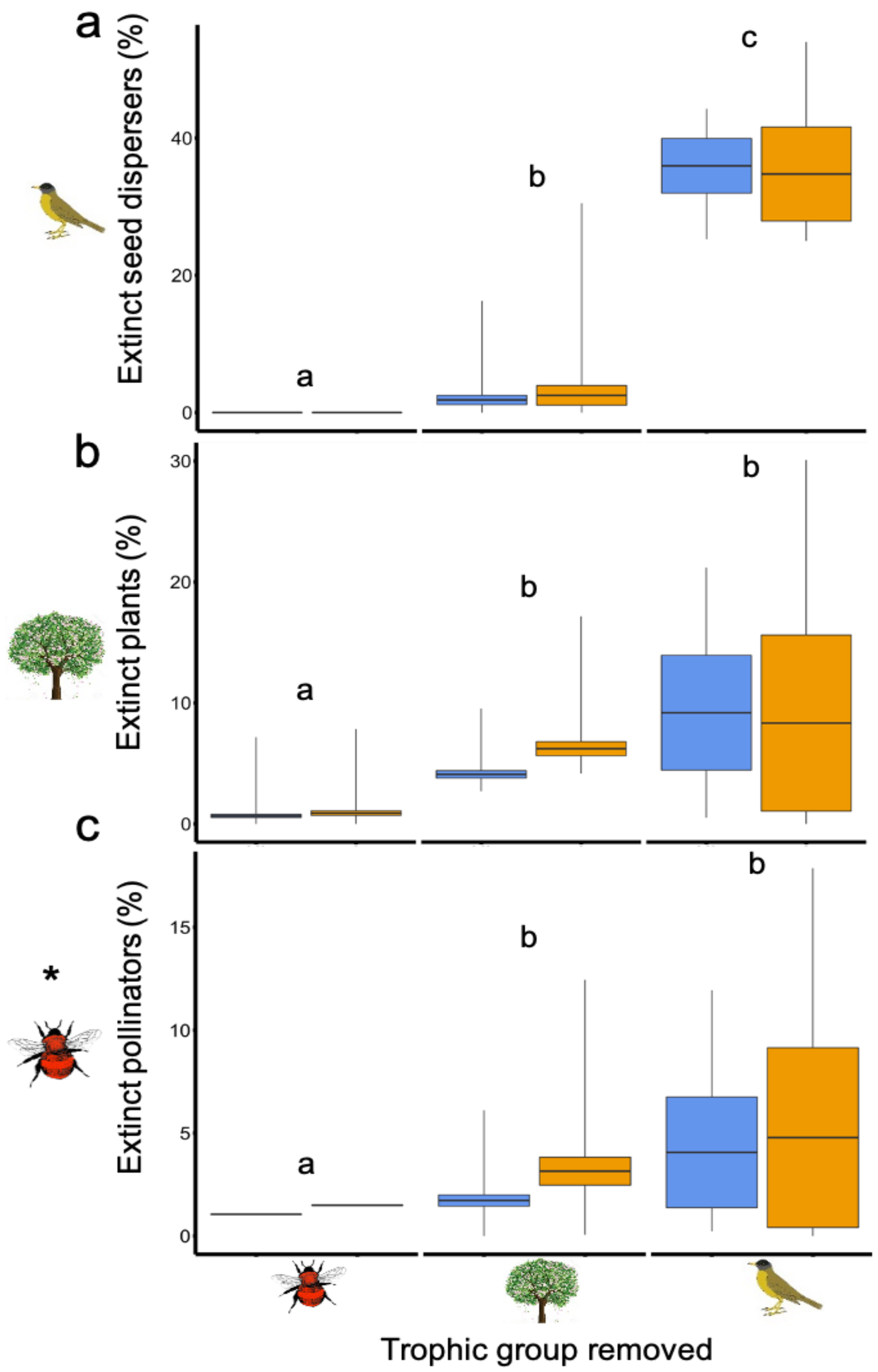


**Figure S15.** Percentage of extinct species in the stochastic coextinction model simulations by tropic group. (a) seed disperser, (b) plant, and (c) pollinators. The % extinct species was calculated at equilibrium following the removal of a single species. Color of boxes indicates types of network: intact (light blue) and invaded (orange). Boxes represent mean ± standard deviation; error bars depict the range. Letters depict significant differences between trophic groups. The asterisk indicates a significant difference in the percentage of extinct pollinators between the intact and invaded network.

**List of species' classification according to their value of dependence on mutualism (R):**

**Table S7.** Classification of plant species according to their dependence on pollination by animals.

| Plant name | Propagation (e.g., vegetative, seed and vegetative, seed) | Self - compatible (yes, no) | Mode of pollination (e.g., wind, animals) | R |
|---|---|---|---|---|
| Aca_ova | seed and vegetative | yes | wind | 0.1 |
| Ade_chi | seed and vegetative | yes | animals | 0.5 |
| Als_aur | seed and vegetative | yes | animals | 0.5 |
| Ari_chi | seed and vegetative | no | animals | 0.6 |
| Aza_mic | seed | yes | animals | 0.9 |
| Bac_sal | seed and vegetative | no | animals | 0.6 |
| Ber_dar | seed and vegetative | no | animals | 0.6 |
| Ber_mic | seed and vegetative | no | animals | 0.6 |
| Cir_vul | seed | yes | animals | 0.9 |
| Cod_les | seed and vegetative | no | animals | 0.6 |
| Col_hys | seed | no | animals | 1 |
| Con_aja | seed | yes | animals | 0.9 |
| Cyn_die | seed | yes | animals | 0.9 |
| Cyt_sco | seed | no | animals | 1 |
| Das_dia | seed and vegetative | yes | animals | 0.5 |
| Emb_coc | seed and vegetative | no | animals | 0.6 |
| Fuc_mag | seed and vegetative | yes | animals | 0.5 |
| Gal_hyp | seed | yes | animals | 0.9 |
| Gau_muc | seed and vegetative | no | animals | 0.6 |
| Gav_odo | seed and vegetative | no | animals | 0.6 |
| Ger_mag | seed | no | animals | 1 |

| | | | | |
|---|---|---|---|---|
| Lab_ana | seed and vegetative | yes | animals | 0.5 |
| Lum_api | seed and vegetative | no | animals | 0.6 |
| May_boa | seed | no | animals | 1 |
| May_chu | seed | no | animals | 1 |
| Mut_dec | seed and vegetative | yes | animals | 0.5 |
| Mut_spi | seed and vegetative | yes | animals | 0.5 |
| Osm_chi | seed | yes | unknown | 0.8 |
| Pha_sec | seed | yes | animals | 0.9 |
| Pla_lan | seed and vegetative | no | wind | 0.1 |
| Pru_avi | seed and vegetative | no | animals | 0.6 |
| Pru_vul | seed and vegetative | yes | animals | 0.5 |
| Rib_mag | seed and vegetative | unknown | animals | 0.5 |
| Ros_rub | seed and vegetative | yes | animals | 0.5 |
| Rub_ida | seed and vegetative | yes | animals | 0.5 |
| Sch_pat | seed | no | animals | 1 |
| Sol_nit | seed and vegetative | yes | animals | 0.5 |
| sp_1 | seed and vegetative | unknown | animals | 0.5 |
| sp_2 | seed and vegetative | unknown | animals | 0.5 |
| sp_3 | seed and vegetative | unknown | animals | 0.5 |
| Tar_offi | seed | yes | animals | 0.9 |
| Tri_cor | seed | yes | animals | 0.9 |
| Tri_pra | seed and vegetative | no | animals | 0.6 |
| Val_sp | seed and vegetative | unknown | animals | 0.5 |
| Vic_nig | seed | yes | animals | 0.9 |
| Vio_mac | seed and vegetative | yes | animals | 0.5 |

Notes: Scientific name of species in online repository.

**Table S2**: Classification of plant species according to their dependence on seed dispersal by animals.

| Plant name | Propagation (e.g. Vegetative, seed and vegetative, seed) | Fleshy-fruited species (yes, no) | R |
|---|---|---|---|
| Aca_ova | seed and vegetative | no | 0 |
| Ade_chi | seed and vegetative | no | 0 |
| Als_aur | seed and vegetative | no | 0 |
| Ari_chi | seed and vegetative | yes | 0.5 |
| Aza_mic | seed | yes | 0.9 |
| Bac_sal | seed and vegetative | no | 0 |
| Ber_dar | seed and vegetative | yes | 0.5 |
| Ber_mic | seed and vegetative | yes | 0.5 |
| Cir_vul | seed | no | 0 |
| Cod_les | seed and vegetative | no | 0 |
| Col_hys | seed | yes | 0.9 |
| Con_aja | seed | no | 0 |
| Cyn_die | seed | no | 0 |
| Cyt_sco | seed | no | 0 |
| Das_dia | seed and vegetative | no | 0 |
| Emb_coc | seed and vegetative | no | 0 |
| Fuc_mag | seed and vegetative | yes | 0.5 |
| Gal_hyp | seed | yes | 0.9 |
| Gau_muc | seed and vegetative | yes | 0.5 |
| Gav_odo | seed and vegetative | no | 0 |
| Ger_mag | seed | no | 0 |
| Lab_ana | seed and vegetative | no | 0 |

| | | | |
|---|---|---|---|
| Lum_api | seed and vegetative | yes | 0.5 |
| May_boa | seed | yes | 0.9 |
| May_chu | seed | yes | 0.9 |
| Mut_dec | seed and vegetative | no | 0 |
| Mut_spi | seed and vegetative | no | 0 |
| Osm_chi | seed | no | 0 |
| Pha_sec | seed | no | 0 |
| Pla_lan | seed and vegetative | No | 0 |
| Pru_avi | seed and vegetative | yes | 0.5 |
| Pru_vul | seed and vegetative | no | 0 |
| Rib_mag | seed and vegetative | yes | 0.5 |
| Ros_rub | seed and vegetative | yes | 0.5 |
| Rub_ida | seed and vegetative | yes | 0.5 |
| Sch_pat | seed | yes | 0.9 |
| Sol_nit | seed and vegetative | yes | 0.5 |
| sp_1 | - | no | 0 |
| sp_2 | - | no | 0 |
| sp_3 | - | no | 0 |
| Tar_offi | seed | no | 0 |
| Tri_cor | seed | yes | 0.9 |
| Tri_pra | seed and vegetative | no | 0 |
| Val_sp | seed and vegetative | no | 0 |
| Vic_nig | seed | no | 0 |
| Vio_mac | seed and vegetative | no | 0 |

Notes: Scientific name of species in online repository.

**Table S3**. Classification of pollinator species according to their dependence on plant resources.

| Pollinator name | R |
|---|---|
| Bom_dah | 0.9 |
| Bom_rud | 0.9 |
| Bom_ter | 0.9 |
| Col_sem | 0.9 |
| Mar_bca | 0.9 |
| Mar_rojmarr | 0.9 |
| mosc_culro | 0.5 |
| Nar_culo_neg | 0.9 |
| Sep_sep | 0.9 |
| Ves_chi | 0.5 |
| Ves_ger | 0.5 |
| sp1 | 0.1 |
| sp2 | 0.5 |
| sp3 | 0.1 |
| sp4 | 0.5 |
| sp5 | 0.1 |
| sp6 | 0.9 |
| sp7 | 0.5 |
| sp8 | 0.1 |
| sp9 | 0.5 |
| sp10 | 0.9 |
| sp11 | 0.9 |
| sp14 | 0.9 |

| | |
|---|---|
| sp15 | 0.1 |
| sp16 | 0.9 |
| sp17 | 0.5 |
| sp18 | 0.9 |
| sp19 | 0.9 |
| sp20 | 0.1 |
| sp21 | 0.5 |
| sp22 | 0.5 |
| sp23 | 0.1 |
| sp25 | 0.1 |
| sp26 | 0.5 |
| sp27 | 0.1 |
| sp28 | 0.9 |
| sp29 | 0.5 |
| sp30 | 0.5 |
| sp31 | 0.5 |
| sp32 | 0.5 |
| sp33 | 0.1 |
| sp34 | 0.5 |
| sp37 | 0.9 |
| sp38 | 0.5 |
| sp39 | 0.5 |
| sp40 | 0.5 |
| sp41 | 0.5 |
| sp42 | 0.9 |
| sp43 | 0.9 |

| | |
|---|---|
| sp44 | 0.5 |
| sp45 | 0.9 |
| sp46 | 0.9 |
| sp47 | 0.9 |
| sp48 | 0.5 |
| sp49 | 0.5 |
| sp50 | 0.1 |
| sp51 | 0.9 |
| sp52 | 0.5 |
| sp53 | 0.5 |
| sp54 | 0.5 |
| sp55 | 0.5 |
| sp56 | 0.1 |
| sp57 | 0.5 |
| sp58 | 0.1 |
| sp59 | 0.5 |
| sp60 | 0.9 |
| sp61 | 0.1 |
| sp62 | 0.5 |
| sp63 | 0.5 |
| sp64 | 0.1 |
| sp65 | 0.5 |
| sp66 | 0.1 |
| sp67 | 0.5 |
| sp70 | 0.1 |
| sp71 | 0.9 |

| | |
|---|---|
| sp72 | 0.5 |
| sp73 | 0.1 |
| sp76 | 0.1 |
| sp77 | 0.9 |
| sp82 | 0.5 |
| sp83 | 0.9 |
| sp84 | 0.9 |
| sp85 | 0.5 |
| sp86 | 0.1 |
| sp87 | 0.1 |
| sp88 | 0.9 |
| sp90 | 0.9 |
| sp91 | 0.9 |
| sp92 | 0.5 |
| sp93 | 0.9 |
| sp94 | 0.9 |
| sp95 | 0.1 |
| sp96 | 0.5 |
| sp97 | 0.9 |
| sp99 | 0.5 |
| sp100 | 0.9 |
| sp101 | 0.9 |
| sp102 | 0.1 |
| sp105 | 0.9 |
| sp106 | 0.5 |
| sp107 | 0.5 |

| | |
|---|---|
| sp108 | 0.5 |
| sp109 | 0.9 |
| sp110 | 0.5 |
| sp111 | 0.9 |
| sp112 | 0.5 |
| sp113 | 0.1 |

Notes: Scientific name of species in online repository.

**Table S4.** Classification of seed disperser species according to their dependence on plant resources. Relevance of each dietary component to species fitness are indicated with the symbols “+”.

| Seed disperser name | Fruits | Invertebrates | Seeds | Eggs | Vertebrates (e.g., birds) | R |
|---|---|---|---|---|---|---|
| Aph_spi | + | +++ | - | - | - | 0.1 |
| Cam_mag | + | +++ | - | - | - | 0.1 |
| Dro_gli | ++ | +++ | - | + | + | 0.5 |
| Ela_alb | +++ | ++ | + | - | - | 0.9 |
| Tur_fal | +++ | ++ | - | - | - | 0.9 |

Notes: Scientific name of species in online repository.

**List of literature reviewed to assign the dependence on mutualism (R) to species:**

Aizen, M. A. (2003). Influences of animal pollination and seed dispersal on winter flowering in a temperate mistletoe. *Ecology*, *84*(10), 2613-2627.

Amico, G. C., & Aizen, M. A. (2005). Dispersión de semillas por aves en un bosque templado de Sudamérica austral:¿ quién dispersa a quién?. *Ecología austral*, *15*(1), 089-100.

Arena, M. E., Lencinas, M. V., & Radice, S. (2018). Variability in floral traits and reproductive success among and within populations of Berberis microphylla G. Forst., an underutilized fruit species. *Scientia Horticulturae*, *241*, 65-73.

Boucher, S., & Pollet, M. (2021). The leaf-miner flies (Diptera: Agromyzidae) of Mitaraka, French Guiana. *Zoosystema*, *43*(6), 113-125.

Bravo, S. P., Cueto, V. R., & Amico, G. C. (2015). Do animal–plant interactions influence the spatial distribution of *Aristotelia chilensis* shrubs in temperate forests of southern South America?. *Plant Ecology*, *216*(3), 383-394.

Caldiz, M. S., & Premoli, A. C. (2006). Isozyme diversity in large and isolated populations of Luma apiculata (Myrtaceae) in north-western Patagonia, Argentina. *Australian journal of botany*, *53*(8), 781-787.

Carvallo, G. O., & Medel, R. (2016). Heterospecific pollen transfer from an exotic plant to native plants: assessing reproductive consequences in an Andean grassland. *Plant Ecology & Diversity*, *9*(2), 151-157.

Celis-Diez, J. L., Hetz, J., Mar'n-Vial, P. A., Fuster, G., Necochea, P., Vásquez, R. A., ... & Armesto, J. J. (2012). Population abundance, natural history, and habitat use by the arboreal marsupial *Dromiciops gliroides* in rural Chiloé Island, Chile. Journal of Mammalogy,93(1), 134-148.

Chang, H., Downie, S. R., Peng, H., & Sun, F. (2019). Floral organogenesis in three members of the tribe Delphinieae (Ranunculaceae). *Plants*, *8*(11), 493.

Cruden, R. W., Baker, K. K., Cullinan, T. E., Disbrow, K. A., Douglas, K. L., Erb, J. D., ... & Wilmot, S. R. (1990). The mating systems and pollination biology of three species of Verbena (Verbenaceae). *Journal of the Iowa Academy of Science: JIAS*, *97*(4), 178-183.

de Lima Ferreira, P., Saavedra, M. M., & Groppo, M. (2019). Phylogeny and circumscription of Dasyphyllum (Asteraceae: Barnadesioideae) based on molecular data with the recognition of a new genus, Archidasyphyllum. *PeerJ*, *7*, e6475

DiTommaso, A., Lawlor, F. M., & Darbyshire, S. J. (2005). The biology of invasive alien plants in Canada. 2. *Cynanchum rossicum* (Kleopow) Borhidi [= *Vincetoxicum rossicum* (Kleopow) Barbar.] and Cynanchum louiseae (L.) Kartesz & Gandhi [= *Vincetoxicum nigrum* (L.) Moench]. *Canadian Journal of Plant Science*, *85*(1), 243-263.

Ducci, F., De Cuyper, B., De Rogatis, A., Dufour, J., & Santi, F. (2013). Wild cherry breeding (Prunus avium L.). In *Forest tree breeding in Europe* (pp. 463-511). Springer, Dordrecht.

Dzendoletas, M. A., Havrylenko, M., & Crivelli, E. (2003). Fenología de plantas en Puerto Blest, Parque Nacional Nahuel Huapi, Patagonia, Argentina. *Ecología*, *17*, 87-98.

Fernández, M. J., López-Calleja, M. V., & Bozinovic, F. (2002). Interplay between the energetics of foraging and thermoregulatory costs in the green-backed firecrown hummingbird *Sephanoides sephaniodes*. *Journal of Zoology*, *258*(3), 319-326.

Fuentealba Jara, I. S. (2016). Germinación in vitro de Orites Myrtoidea (Proteaceae) y *Maytenus Chubutensis* (Celastraceae). Especies vegetales insuficientemente conocidas de la Flora de Chile.

Funk, V. A., Pasini, E., Bonifacino, J. M., & Katinas, L. (2016). Home at last: the enigmatic genera Eriachaenium and Adenocaulon (Compositae, Mutisioideae, Mutisieae, Adenocaulinae). *PhytoKeys*, (60), 1.

Gerhardt, R. R., & Hribar, L. J. (2019). Flies (Diptera). In *Medical and Veterinary Entomology* (pp. 171-190). Academic Press.

Godoy, M., De La Fuente, L. M., Gómez, M., & Ginocchio, R. (2020). Aspectos reproductivos, arquitectura y fenomorfología de Maytenus boaria Molina (Celastraceae) en Chile central. *Gayana. Botánica*, *77*(2), 152-167.

Graça, V. C., Ferreira, I. C., & Santos, P. F. (2020). Bioactivity of the Geranium genus: a comprehensive review. *Current pharmaceutical design*, *26*(16), 1838-1865.

Huryn, V. M. B., & Moller, H. (1995). An assessment of the contribution of honey bees (Apis mellifera) to weed reproduction in New Zealand protected natural areas. *New Zealand Journal of Ecology*, 111-122.

Hyslop, M. G., Kemp, P. D., & Hodgson, J. (1999, January). Vegetatively reproductive red clovers (*Trifolium pratense* L.): an overview. In *Proceedings of the New Zealand Grassland Association* (pp. 121-126).

Karolyi, F., Szucsich, N. U., Colville, J. F., & Krenn, H. W. (2012). Adaptations for nectar-feeding in the mouthparts of long-proboscid flies (Nemestrinidae: Prosoeca). *Biological Journal of the Linnean Society*, *107*(2), 414-424.

Klinkhamer, P. G., & De Jong, T. J. (1993). Cirsium vulgare (Savi) Ten.(Carduus lanceolatus L., Cirsium lanceolatum (L.) Scop., non Hill). *Journal of Ecology (Oxford)*, *81*(1), 177-191.

Kutschker, A. (2011). Revisión del género Valeriana (Valerianaceae) en Sudamérica austral. *Gayana. Botánica*, *68*(2), 244-296.

Ladio, A. H., & Aizen, M. A. (1999). Early reproductive failure increases nectar production and pollination success of late flowers in south Andean *Alstroemeria aurea*. *Oecologia*, *120*(2), 235-241.

Lamas, C. J. E., & Evenhuis, N. L. (2016). Family Bombyliidae. *Catalogue of Diptera of Colombia*, *4122*(1), 372-381.

Lawrence, J. F., & Leschen, R. A. (2011). 9.11. Melyridae Leach, 1815. In *Morphology and Systematics (Elateroidea, Bostrichiformia, Cucujiformia partim)* (pp. 273-280). De Gruyter.

Ling, T. C., Wang, L. L., Zhang, Z. Q., Dafni, A., Duan, Y. W., & Yang, Y. P. (2017). High autonomous selfing capacity and low flower visitation rates in a subalpine population of *Prunella vulgaris* (Lamiaceae). *Plant Ecology and Evolution*, *150*(1), 59-66.

Lowry, P., & Jones, A. (1984). Systematics of Osmorhiza Raf. (Apiaceae: Apioideae). *Annals of the Missouri Botanical Garden, 71*(4), 1128-1171. doi:10.2307/2399249

Marticorena, A. E., & Cavieres, L. A.. (2000). *Acaena magellanica* (Lam.) Vahl (Rosaceae). *Gayana. Botánica*, *57*(1), 107-113. https://dx.doi.org/10.4067/S0717-66432000000100011

Medan, D., & Torretta, J. P. (2015). The reproduction of Colletia hystrix and late-flowering in Colletia (Rhamnaceae: Colletieae). *Plant Systematics and Evolution*, *301*(4), 1181-1189.

Misle, E., Garrido, E., Contardo, H., & González, W. (2011). Maqui (Aristotelia chilensis (Mol.) Stuntz) the amazing chilean tree: a review. *Journal of Agricultural Science and Technology B*, *1*, 473-482.

Muñoz, C. E., Ippi, S. G., Celis Diez, J. L., Salinas, D., & Armesto, J. J. (2017). Arthropods in the diet of the bird assemblage from a forested rural landscape in Northern Chiloé Island, Chile: a quantitative study.

Narendran, T. C., & Rao, S. A. (1987). Biosystematics of Chalcididae (Chalcidoidea: Hymenoptera). *Proceedings: Animal Sciences*, *96*(5), 543-550.

Orellana, J. I., Smith-Ramírez, C., Rau, J. R., Sade, S., Gantz, A., & Valdivia, C. E. (2014). Phenological synchrony between the austral thrush *Turdus falcklandii* (Passeriformes: Turdidae) and its food resources within forests and prairies in southern Chile. *Revista chilena de historia natural*, *87*(1), 1-8.

Powell, K. I., Krakos, K. N., & Knight, T. M. (2011). Comparing the reproductive success and pollination biology of an invasive plant to its rare and common native congeners: a case study in the genus Cirsium (Asteraceae). *Biological Invasions*, *13*(4), 905-917.

Romoleroux, K., Cárate-Tandalla, D., Erler, R., Navarrete, H. 2019. *Galium hypocarpium* En: Plantas vasculares de los bosques de *Polylepis* en los páramos de Oyacachi. Version 2019.0 https://bioweb.bio/floraweb/polylepis/FichaEspecie/Galium%20hypocarpium

Rosenberger, N. M. (2018). Competition of a nectar-robbing bumble bee with a legitimate forager and its consequences for female reproductive success of Fuchsia magellanica (Unpublished master's thesis). University of Calgary, Calgary, AB. doi:10.11575/PRISM/33042 http://hdl.handle.net/1880/108689 master thesis

Saavedra Cárdenas, M. J. (2016). Adaptación de protocolos de establecimiento in vitro de *Ribes rubrum* L., *Ribes nigrum* L. y *Ribes uva-crispa* L.

Seguí, J., Lázaro, A., Traveset, A., Salgado-Luarte, C., & Gianoli, E. (2018). Phenotypic and reproductive responses of an Andean violet to environmental variation across an elevational gradient. *Alpine Botany*, *128*(1), 59-69.

Sharma, N., Koul, P., & Koul, A. K. (1993). Pollination biology of some species of genus Plantago L. *Botanical Journal of the Linnean Society*, *111*(2), 129-138.

Silva, V. C., & Mello, R. L. (2008). Occurrence of *Physoclypeus farinosus* Hendel (Diptera: Lauxaniidae) in Flowerheads of Asteraceae (Asterales). *Neotropical entomology*, *37*, 92-96.

Smith-Ramírez, C., Martinez, P., Nunez, M., González, C., & Armesto, J. J. (2005). Diversity, flower visitation frequency and generalism of pollinators in temperate rain forests of Chiloé Island, Chile. *Botanical Journal of the Linnean Society*, *147*(4), 399-416.

Soza, V. L., & Olmstead, R. G. (2010). Evolution of breeding systems and fruits in New World Galium and relatives (Rubiaceae). *American Journal of Botany*, *97*(10), 1630-1646.

Stawiarz, E., & Wróblewska, A. (2013). Flowering dynamics and pollen production of *Laburnum anagyroides* Med. under the conditions of South-Eastern Poland. *Journal of Apicultural Science*, *57*(2), 103.

Suzuki, N. (2003). Significance of flower exploding pollination on the reproduction of the Scotch broom, *Cytisus scoparius* (Leguminosae). *Ecological research*, *18*(5), 523-532

Teillier, S., & Escobar, F. (2013). Revisión del género Gaultheria L.(Ericaceae) en Chile. *Gayana. Botánica*, *70*(1), 136-153.

Troiani, H. O. (1985). Las especies de Baccharis (Compositae) de la provincia de La Pampa. *Revista Facultad de Agronomía Universidad Nacional de la Pampa*, *1*, 1-2.

Ueda, Y., & Akimoto, S. (2001). Cross-and self-compatibility in various species of the genus Rosa. *The Journal of Horticultural Science and Biotechnology*, *76*(4), 392-395.

Vallejo-Marín, M., Walker, C., Friston-Reilly, P., Solís-Montero, L., & Igic, B. (2014). Recurrent modification of floral morphology in heterantherous Solanum reveals a parallel shift in reproductive strategy. *Philosophical Transactions of the Royal Society B: Biological Sciences*, *369*(1649), 20130256.

Vázquez, D. P., & Simberloff, D. (2004). Indirect effects of an introduced ungulate on pollination and plant reproduction. *Ecological Monographs*, *74*(2), 281-308.

Vergara, P., & Schlatter, R. P. (2004). Magellanic Woodpecker (*Campephilus magellanicus*) abundance and foraging in Tierra del Fuego, Chile. *Journal of Ornithology*, *145*(4), 343-351.

Wilson, J. S., & Carril, O. J. M. (2015). 4. COLLETIDAE. In *The Bees in Your Backyard* (pp. 96-110). Princeton University Press.

Żurawicz, E. (2015). Cross-pollination increases the number of drupelets in the fruits of red raspberry *(Rubus idaeus* L.). In *XI International Rubus and Ribes Symposium 1133* (pp. 145-152).
